# VMAT2 dysfunction impairs vesicular dopamine uptake, driving its oxidation and α-synuclein pathology in DJ-1-linked Parkinson’s disease neurons

**DOI:** 10.1101/2025.09.22.677565

**Authors:** Leonie M. Heger, Francesco Gubinelli, Andreas Huber, Aida Cardona-Alberich, Matteo Rovere, Ulf Matti, Stephan A. Müller, Sankarshana R. Nagaraja, Lena Jaschkowitz, Martina Schifferer, Wolfgang Wurst, Stefan F. Lichtenthaler, Christian Behrends, Sivakumar Sambandan, Lena F. Burbulla

**Affiliations:** Metabolic Biochemistry, Biomedical Center (BMC), Faculty of Medicine, LMU Munich, Munich, Germany; Abberior Instruments GmbH, Göttingen, Germany; Neuroproteomics, School of Medicine and Health, TUM University Hospital, Technical University of Munich, Germany; German Center for Neurodegenerative Diseases (DZNE), Munich, Germany; 5Munich Cluster for Systems Neurology (SyNergy), Munich, Germany; Faculty of Medicine, LMU Munich, Munich, Germany; Helmholtz Munich, Institute of Developmental Genetics, Munich, Germany, Technical University Munich, Institute of Developmental Genetics, Munich, Germany; Laboratory of Neurobiology, Max Planck Institute for Multidisciplinary Sciences, Göttingen, Germany; Synaptic Metal Ion Dynamics and Signaling, Max Planck Institute for Multidisciplinary Sciences, Göttingen, Germany

**Author notes:** Corresponding author: Lena Burbulla, PhD, Professor for Metabolic Biochemistry of Neurodegenerative Diseases, Biomedical Center (BMC), Faculty of Medicine, Ludwig-Maximilians-Universität München & German Center for Neurodegenerative Diseases (DZNE) Munich.

## Abstract

Parkinson’s disease (PD) is characterized by α-synuclein accumulation and dopaminergic neuron degeneration, with dopamine (DA) oxidation emerging as a key pathological driver. However, the mechanisms underlying this neurotoxic process remain unclear. Using PD patient-derived and CRISPR-engineered iPSC midbrain dopaminergic neurons lacking DJ-1, we identified defective sequestration of cytosolic DA into synaptic vesicles, which culminated in DA oxidation and α-synuclein accumulation. In-depth proteomics, state-of-the-art imaging, and ultrasensitive DA probes uncovered that decreased VMAT2 protein and function impaired vesicular DA uptake, resulting in reduced vesicle availability and abnormal vesicle morphology.

Furthermore, VMAT2 activity and vesicle endocytosis are processes dependent on ATP, which is notably reduced in DJ-1-deficient dopaminergic neurons. ATP supplementation restored vesicular function and alleviated DA-related pathologies in mutant dopaminergic neurons.

This study reveals an ATP-sensitive mechanism that regulates DA homeostasis through VMAT2 and vesicle dynamics in midbrain dopaminergic neurons, highlighting enhanced DA sequestration as a promising therapeutic strategy for PD.

**Teaser:** Loss of DJ-1 interferes with VMAT2 function and vesicle dynamics, leading to DA oxidation and α-synuclein pathology in PD neurons.

## Introduction

Understanding the pathophysiology of neurodegenerative diseases is crucial for identifying new therapeutic targets and developing effective treatments. Parkinson’s disease (PD), the second most common neurodegenerative disorder, affects approximately 1% of individuals over 65 and 4% of those over 85. While the majority of cases are sporadic, 5%–10% are attributed to inherited cases (*1*). PD is marked by the pathological accumulation of α-synuclein into Lewy bodies and the progressive loss of dopaminergic neurons in the substantia nigra (SN), leading to characteristic motor symptoms such as tremor and rigidity (*2*). Current treatments offer only symptomatic relief without slowing neuronal degeneration. Our understanding of the pathways underlying the differential vulnerability of SN dopaminergic neurons in PD remains limited. Uncovering the mechanisms behind the depletion of this specific neuron subpopulation and understanding the progression of disease in more detail is crucial for the development of disease-modifying therapies.

A time-dependent pathological cascade has been recently uncovered, starting with mitochondrial dysfunction and leading to dopamine (DA) oxidation and α-synuclein accumulation in DJ-1 (*PARK7*)-linked PD as well as in other genetic and sporadic forms of PD (*3*). Notably, this pathogenic cascade was not recapitulated in PD mouse models, and species-specific differences in DA metabolism are likely to contribute to this discrepancy. In particular, PD mouse models lacked the accumulation of oxidized DA characteristic of human midbrain neurons (*3*). This fundamental difference suggests that as-yet-unknown mechanisms prevent the formation of oxidized DA in mice. This may, at least in part, explain the relative resistance of rodent neurons to degeneration in genetic models of PD, while also underscoring the critical role of DA oxidation in the vulnerability of human SN neurons to PD-linked mutations.

In general, the oxidation of DA is dependent on various factors, beginning with an excess of cytosolic DA that is not properly sequestered into synaptic vesicles (*4*). If, for various reasons, this pathway fails or is reaching its capacity, DA is enzymatically degraded through specific enzymes (*5*). This sensitive regulation is particularly important as the accumulation of cytosolic DA generates reactive oxygen species (ROS) and DA quinones (DAQs) (*6, 7*), derivates of oxidized DA, that can form toxic adducts with proteins (*3, 8, 9*), and may lead to neuron damage and death (*10*). This pathway finally results in the formation of neuromelanin (NM), a polymer pigment concentrated in human – but not mouse – SN midbrain DA neurons (*11*). NM is thought to play a protective role by preventing toxic DA accumulation and scavenging metals (*12*). However, while altered mitochondrial function was suggested to contribute to oxidized DA formation and broader PD pathogenesis, the mechanisms by which mitochondrial dysfunction leads to excess cytosolic DA, triggering its oxidation, and whether targeting mitochondrial health would work as therapeutic approach to counteract oxidized DA is largely unexplored. Dopaminergic neurons are particularly vulnerable to oxidative stress due to their autonomous pacemaking activity and extensive axonal arborization, both of which contribute to elevated mitochondrial oxidant stress (*13*).

It has been well established that DA signaling and metabolism are largely regulated by the synergistic interplay of tyrosine hydroxylase (TH), vesicular monoamine transporter 2 (VMAT2), and DA transporter (DAT). While TH catalyzes the conversion of tyrosine to L-3,4-dihydroxyphenylalanine (L-DOPA), which is subsequently converted to DA by DOPA decarboxylase, VMAT2 and DAT are specific membrane transporters responsible for transporting of DA from the cytosol into synaptic vesicles and across the plasma membrane into the extracellular space, respectively. In particular, VMAT2 requires a proton electrochemical gradient generated in the synaptic vesicle lumen by vacuolar ATPase, which, in turn, depends on ATP primarily sourced from local mitochondria, to package DA (*14*) from the cytosol into synaptic vesicles (*15, 16*). Therefore, VMAT2 relies on a well-working mitochondrial function for sufficient ATP delivery. By regulating the concentration or number of DA neurotransmitter molecules sequestered per vesicle, i.e. the quantal size, VMAT2 also largely determines the quantity of DA release into the synaptic cleft (*15*). Interestingly, studies have indicated a reduction in VMAT2 activity and DA uptake in isolated DA storage vesicles from striatal PD brain tissue (*17*). VMAT2 dysfunction is therefore one potential source for toxic DA products due to increased unsequestered cytosolic DA. However, why VMAT2 activity is reduced in PD brain tissue – and whether targeting VMAT2 deficiency or improving its function could be beneficial in human disease models or even therapeutically – remain poorly investigated, particularly in disease-relevant human model systems.

In this study, we aimed at investigating underlying mechanisms of DA neuron vulnerability linked to factors leading to oxidized DA accumulation. Using induced pluripotent stem cell (iPSC)-derived midbrain dopaminergic neurons from a DJ-1-linked PD patient and two *DJ-1* knock-out (KO) lines, our results strongly link the loss of this redox-sensitive protein with a reduction in VMAT2 abundance across multiple magnitudes of spatial dimensions. In addition to this, VMAT2 transporter activity was diminished due to lowered ATP levels in DJ-1-deficient neurons. These combined pathologies resulted in a reduced uptake of DA into synaptic vesicles leading to oxidized DA and α-synuclein accumulation in mutant neurons. Unbiased and targeted proteomic analyses strengthened the link of DJ-1 with intracellular vesicles and synaptic components, an association that is heavily diminished in DJ-1-deficient neurons. Finally, we show that targeting ATP depletion in these PD-linked neurons partially alleviates the observed PD-associated pathologies, offering a potential approach benefiting neuronal survival. Our study provides insights into whether and how the main route of DA sequestration through VMAT2-mediated uptake into synaptic vesicles is altered in PD-linked neurons, a potential underlying mechanism for DA oxidation and vulnerability of SN midbrain neurons.

## Results

### Proteomic analysis reveals deficiency of VMAT2 and downregulation of pathways associated with synaptic vesicles in DJ-1-deficient dopaminergic neurons

Oxidized DA accumulation has been described in DJ-1-deficient iPSC-derived midbrain dopaminergic neurons (*3*), however, the source of this potentially neurotoxic insult affecting neuron vulnerability remains elusive. To uncover underlying pathways of the observed DA oxidation, we differentiated iPSCs from two *PARK7*/*DJ-1* KO lines (*DJ-1* KO #1 and #2 = DJ-1^KOs^) generated by gene-editing using clustered regularly interspaced short palindromic repeats (CRISPR)–Cas9 and their respective healthy control counterparts (CTRL KO #1 and #2 = CTRLs) (*3*) into midbrain dopaminergic neurons (fig. S1 and S2).

In detail, iPSCs expressed key pluripotency markers (SSEA4, OCT4, NANOG, SOX2) as confirmed by immunocytochemistry (fig. S1A to B). qPCR analysis showed strong upregulation of pluripotency-related genes compared to human fibroblasts (fig. S1C). Spontaneous differentiation demonstrated the capacity of all lines to generate cells from all three germ layers (ectoderm, mesoderm, endoderm), as shown by qPCR (fig. S1D). Quality control of dopaminergic neurons at days 30 and 70 of differentiation (fig. S2A to B) revealed homogeneous, sustained expression of dopaminergic and neuronal markers, indicating successful induction and maturation. Western blot analysis confirmed expression of TH and β-III-tubulin and validated the loss of DJ-1 in both *DJ-1* KO lines (fig. S2C).

Using these two pairs of CRISPR-edited *DJ-1* KO neuronal lines with their respective isogenic counterparts, we performed unbiased mass spectrometry (MS)-based proteomics. Here, we identified two distinct sets of proteins that are upregulated (385 proteins) and downregulated (104 proteins) in neurons from both *DJ-1* KO lines (p-value < 0.05 and log2 fold change larger than 0.5 or smaller than -0.5) compared to controls (Fig. 1A and table S1). Most interestingly, VMAT2 was discovered among the top three downregulated proteins in the dataset, following PARK7 (DJ-1) as the most downregulated, and PCDAC (also known as PCDHA12, Protocadherin Alpha-12), a protocadherin family member functioning as a calcium-dependent cell-adhesion molecule, as the second. This emphasizes a pronounced role for VMAT2 deficiency in *DJ-1* KO neurons. Other proteins found to be differentially regulated in *DJ-1* KO neurons further indicated alterations in synaptic function in association with neuronal signaling, such as the decreased abundance of PTPRT (Protein Tyrosine Phosphatase Receptor Type T), identified as risk factor common to sporadic PD (*18, 19*), and AT2A3 (also known as SERCA3, Sarco/Endoplasmic Reticulum Ca²⁺-ATPase 3) (*20*). While the direct involvement of AT2A3 in PD is not well studied, its role in pumping calcium from the cytosol in the endoplasmic reticulum and by this regulating calcium homeostasis makes AT2A3 a potential player in the PD-associated pathways.

**Figure 1:**
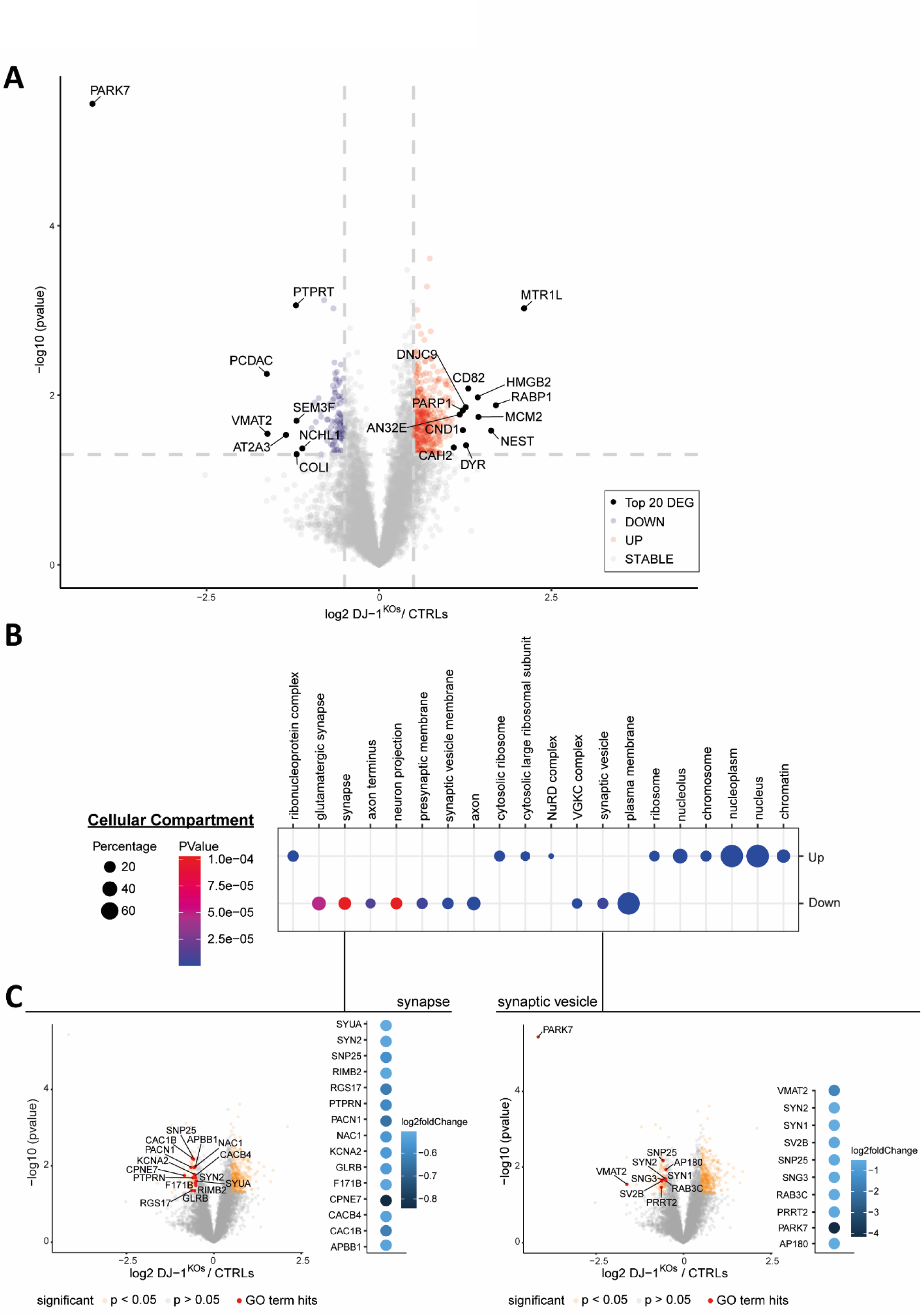
Whole-cell proteomic analysis of DJ-1-deficient dopaminergic neurons. (**A**) Volcano plots of *DJ-1* KO (both lines) versus isogenic control (both lines) dopaminergic neurons. The -log10 transformed p-values of each protein are plotted against their log2 fold change transformed protein label free quantification ratios. Proteins significantly more abundant in the proteome of *DJ-1* KO neurons are displayed as red filled dots, while less abundant proteins are displayed as blue filled dots. Unaltered proteins are displayed as gray dots. Top 20 differentially-expressed proteins are labeled with their UniProt gene names (n = 4). (**B**) Results of gene ontology (GO) enrichment analysis for cellular component (GOTERM_CC_DIRECT) using the web tool DAVID (v 6.8). Differentially expressed proteins (p-value < 0.05 and log2 fold change > 0.5 or < -0.5) were compared to the background of all proteins detected in both conditions. The dot plots show the total gene number of a term in percentage as dot size and the p-values as a color gradient. The top 10 terms for either enriched (Up) or downregulated (Down) proteins are presented. (**C**) Volcano and dot plots of the differentially expressed proteins in *DJ-1* KO neurons, involved in the terms “synapse” (left) and “synaptic vesicle” (right) associated with cellular compartment.

In addition, we performed Gene Ontology (GO) enrichment analysis for cellular component (Fig. 1B), molecular function (fig. S3A), and biological process (fig. S4A) of the differentially expressed proteins. We ranked the top 10 GO terms for either up- or downregulated proteins by significance and, among others, discovered a prominent enrichment of downregulated proteins for the terms “synapse” and “synaptic vesicle” associated with cellular compartment (Fig. 1B), “transmembrane transporter binding” and “calcium ion binding” associated with molecular function (fig. S3A), as well as “chemical synaptic transmission” associated with biological process (fig. S4A) in *DJ-1* KO neurons.

We performed a more in-depth analysis of the above outlined downregulated pathways and visualized the respective differentially expressed proteins of these pathways via volcano plots and dot plots (Fig. 1C, fig. S3B and S4B). Importantly, VMAT2 prominently presented with the second highest fold change in the pathway “synaptic vesicle” (following the top hit *PARK7*, DJ-1) and the highest fold change in “chemical synaptic transmission”.

The aforementioned AT2A3 displayed the highest fold change among proteins in the pathway “transmembrane transporter binding” (fig. S3B). In addition, numerous other proteins represented in these pathways have crucial roles for synaptic function and proper vesicle formation. Out of those, examples are SV2B (synaptic versicle glycoprotein 2B) that is located on synaptic vesicles and involved in neurotransmitter release regulation (*21*) and AT1A1 (alpha-1A subunit of the Na⁺/K⁺-ATPase), an enzyme essential for maintaining the electrochemical gradients across the plasma membrane. Reduced AT1A1 has been linked to α-synuclein pathology in PD (*22*). Furthermore, SYN1 (synapsin 1) and SYN2 (synapsin 2), primarily involved in the regulation of neurotransmitter release and synaptic vesicle dynamics (*23, 24*), were found to be reduced in our cultured *DJ-1* KO neurons. Other proteins essential for clathrin-mediated endocytosis, synaptic vesicle recycling, vesicle dynamics and neurotransmitter release, such as PACN1, RIMB2, RAB3C, or SNP25 (*25–27*), were present in low abundance in *DJ-1* KO neurons. Beyond alterations in proteins involved in DA uptake, synaptic vesicle dynamics, and neurotransmitter release, *DJ-1* KO neurons also showed impaired calcium handling, including reduced expression of CAC1B (also known as CACNA1B), which encodes the Cav2.2 subunit of N-type voltage-gated calcium channels (fig. S3B). This suggests a previously underappreciated role for DJ-1 in presynaptic calcium regulation.

Collectively, our findings highlight VMAT2 to be significantly diminished in DJ-1-deficient neurons together with a substantial disruption of synapse-associated pathways that highlights a potential risk for proper uptake of DA from the cytosol, potentially causing it to oxidize and may, at least partly, explain why this type of neurons is particularly vulnerable.

### DJ-1 interacts with vesicular and cytoskeletal components – an association lost in DJ-1-deficient dopaminergic neurons

Since we found the loss of DJ-1 to affect the abundance of VMAT2 and interfere with pathways associated with synaptic vesicles, we sought to substantiate this notion with a comparative interactome analysis of DJ-1 via co-migration and immunoprecipitation (IP) studies. Therefore, we subjected digitonin-soluble (1%) lysates of *DJ-1* KO and isogenic control neurons to non-denaturing blue-native PAGE (BN-PAGE) to visualize and separate multi-subunit protein complexes associated with DJ-1 (Fig. 2A). When blotting against DJ- 1 after BN-PAGE, we identified a range of native molecular weights with the strongest immunoreactivity in the control neurons physiologically expressing wild-type (WT) DJ-1, and excised four gel blocks (A-D) covering this region, which were then processed via in-gel digestion and liquid chromatography–tandem mass spectrometry (LC-MS/MS) (Fig. 2B and table S2). By comparing the proteomic profiles of control and *DJ-1* KO lysates in each of these bands, we sought to identify potential interactants co-migrating with DJ-1 (Fig. 2C and fig. S5A). Through this approach, proteins and protein complexes coincidentally co-migrating – but not interacting – with DJ-1 would not be differentially enriched, as we would expect these species to be equally abundant in *DJ-1* KO and control lysates. GO cellular component analysis, using SubcellulaRVis (*28*), of the putative interactants co-migrating with DJ-1, unraveled, across all blocks, cytoplasmic and nuclear components enriched in control over *DJ-1* KO lysates (Fig. 2D and fig. S5B). Intriguingly, in bands “A” and “B” we uncovered a significant number of proteins co-migrating with DJ-1 associated with intracellular vesicle compartments (block “A”) and the cytoskeleton (blocks “A”, “B”, “D”) (Fig. 2D and fig. S5B). Peroxisomal proteins were also found to associate with DJ-1 (block “D”, fig. S5B), besides an expected pool of mitochondrial proteins (Blocks “B” and “C”, Fig. 2D and fig. S5B).

**Figure 2:**
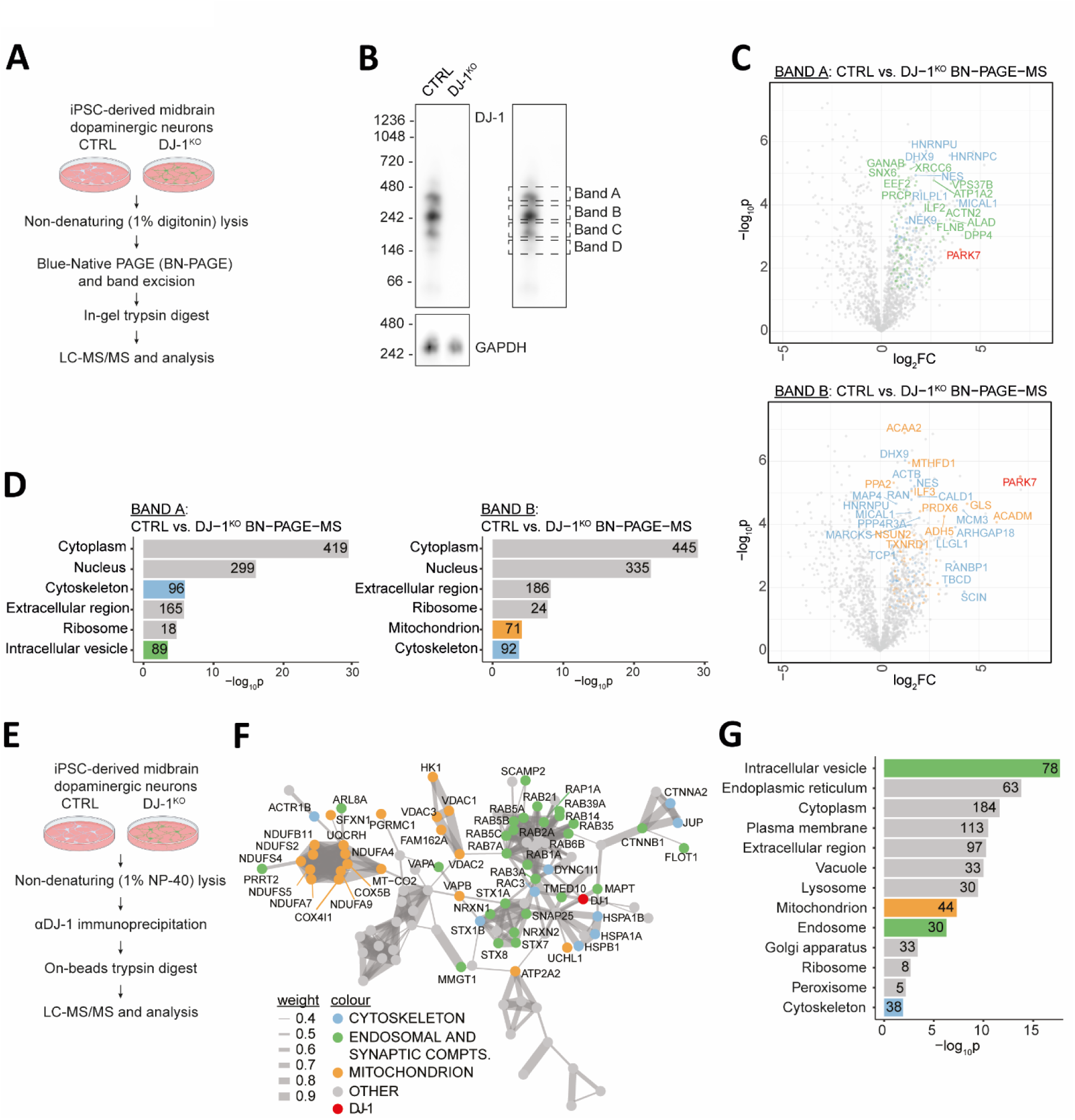
Vesicular association of DJ-1 as defined by co-migration on BN-PAGE and IP-MS. (**A**) Workflow of the co-migration experiment (BN-PAGE followed by LC-MS/MS). (**B**) Lysates from control or *DJ-1* KO neurons were separated on BN-PAGE and four gel bands, covering the region with the strongest DJ-1 immunoreactivity, were excised for LC-MS/MS analysis (bands “A”, “B”, “C”, “D”). (**C**) Volcano plots of the proteins in bands “A” and “B” co-migrating with DJ-1, identified by comparing control to *DJ-1* KO neurons (two-sided t-test, n = 4 biological replicates). Highlighted are representative DJ-1 co-migrants associated with the cytoskeleton (blue), intracellular vesicles (green), and mitochondria (orange). PARK7 (DJ-1) is highlighted in red. (**D**) GO cellular component analysis, using SubcellulaRVis, of DJ-1 co-migrants in bands “A” and “B” (log2FC > 0.5, p < 0.05). (**E**) Workflow of the DJ-1 IP-MS experiment. (**F**) STRING enrichment analysis of DJ-1 co-immunoprecipitated (l og2FC > 2 in 2 out of the 3 biological replicates) proteins. Only experiment- and database-derived physical interactions (confidence ≥ 0.4) were considered to generate the network and only the largest connected sub-network is shown. Color-coding as in (C). (**G**) GO cellular component analysis, using SubcellulaRVis, of DJ-1 co-immunoprecipitated proteins (log2FC > 2 in 2 out of the 3 biological replicates).

Since our co-migration studies supported the association of DJ-1 with vesicular compartments and vesicle-trafficking machinery, we validated this hypothesis by confirming the physical interaction of DJ-1 with vesicular and endosomal protein components via IP-MS of *DJ-1* KO and isogenic control neuronal lysates, using endogenous DJ-1 as our bait (Fig. 2E). On the 221 co-IP’d proteins (including isoforms not resolved by MS, table S3), we ran a STRING enrichment meta-analysis (*29*), restricting our search to established physical interactions. STRING connected a large number of co-IP’d proteins to a single network, suggesting that DJ-1 associates with known protein machineries of synaptic and mitochondrial components, and highlighting, among these, an enrichment of small Rab GTPases and respiratory chain proteins (Fig. 2F). Of note, cytoplasmic and nuclear components, the top GO hits in our co-migration analysis, are much less represented in the co-IPs, where more stringent lysis and wash conditions are applied, suggesting that most of these interactions are very weak and/or spurious. Consistently, subsequent GO cellular component analysis of the co-IP’d interaction partners of DJ-1 showed the strongest enrichment for proteins associated with intracellular vesicles, but also included mitochondria, endosomes, and the cytoskeleton (Fig. 2G). Taken together, MS analysis after BN-PAGE and co-IP strengthen our whole cell proteomics results and confirm the association of DJ-1 with vesicular components in midbrain dopaminergic neurons, an association that is lost upon DJ-1 deficiency.

### Ultrastructural deficiency of VMAT2 synapses in DJ-1-deficient dopaminergic neurons

We wanted to further disentangle the loss of VMAT2 protein in DJ-1-deficient dopaminergic neurons on multiple spatial scales using state-of-the-art imaging approaches. First, confocal microscopy analysis revealed a reduced density of VMAT2-positive synapses in *DJ-1* KO neurons compared to the isogenic healthy control neurons (Fig. 3A) while there was no change in the abundance of VGAT-positive and VGLUT2-positive synapses that were detected in low amounts in this culture (fig. S6). While the molecular and ultrastructural organization of VGAT- and VGLUT2-positive synapses have been extensively studied, dopaminergic synapses are relatively understudied and the molecular composition and structural signatures are poorly understood.

**Figure 3:**
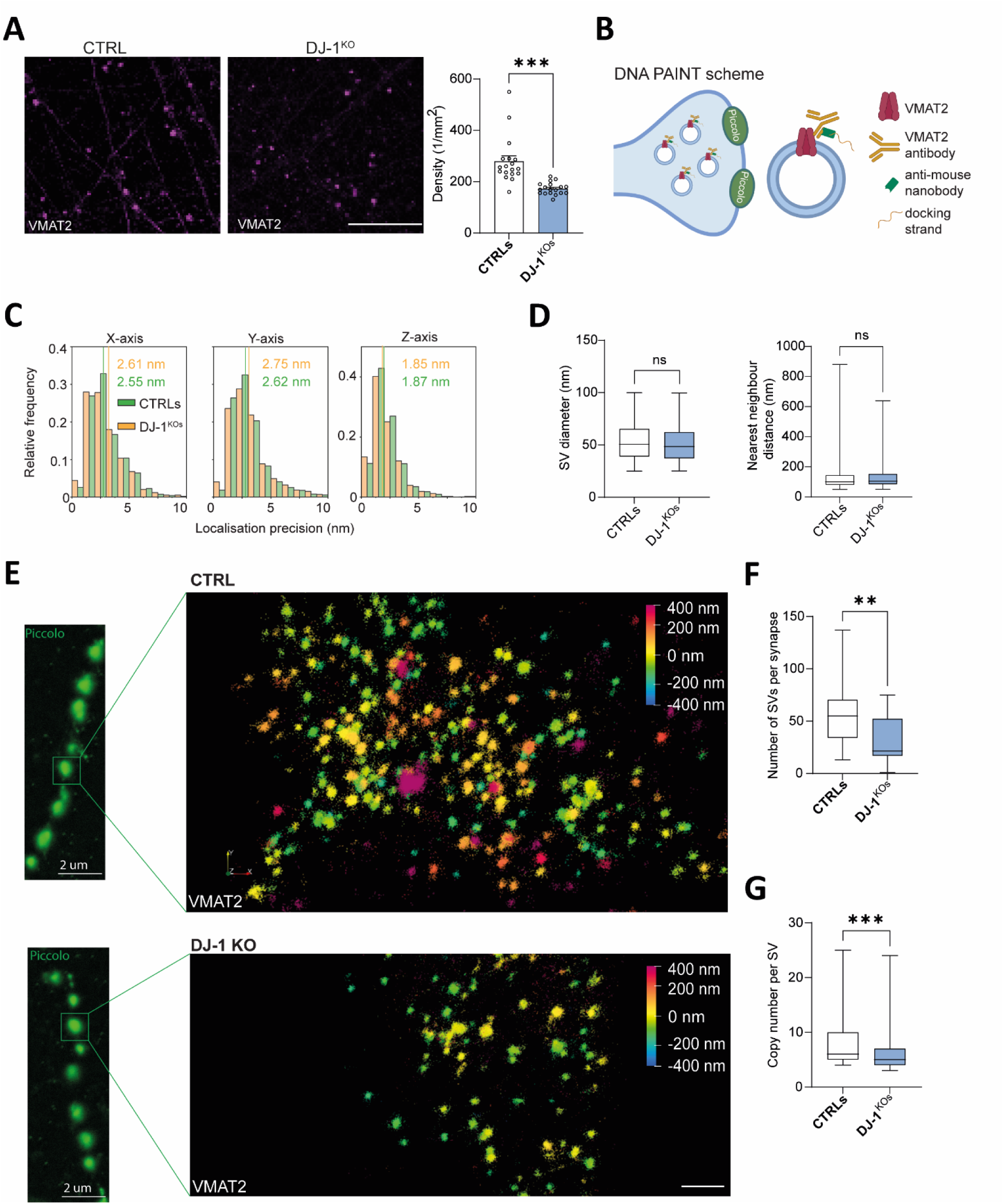
Ultrastructural deficiency of VMAT2 synapses in DJ-1-deficient dopaminergic neurons. (**A**) Representative confocal images of dopaminergic synapses, visualized by staining VMAT2, in control and *DJ-1* KO neurons. Right, bar graph quantifying the density of synapses per area. Scale bar, 20µm (n=18). (**B**) Illustration showing Minflux-DNA PAINT labeling of VMAT2-positive synapses. Piccolo, the active zone marker, was used to identify synapses prior to Minflux imaging. (**C**) Histograms showing the distribution of localization precision for VMAT2 Minflux imaging in the axial (X and Y) and lateral (Z) axis. The yellow and green lines indicate the median for control and *DJ-1* KO neurons, respectively. (**D**) Box-whisker plot quantification of the synaptic vesicle (SV) diameter and the nearest neighbor distance (NND) between vesicles in control and *DJ-1* KO neurons (n=459 vs 849 for SV diameter; n=594 vs 1193 for NND). (**E**) Left, representative confocal image of a dendritic segment showing individual synapses stained by Piccolo. Scale bar, 2µm. Right, Representative Minflux raw images of the zoomed-in region from the left panel showing 3D localizations of VMAT2 in a control (top panel), and a *DJ-1* KO neuron (bottom panel). The individual localizations are color coded by their positions in the z-axis. XY-Scale bar, 100nm. Z-Scale bar, 800nm. (**F**) Box-whisker plot quantification of the number of synaptic vesicles per synapse in control and *DJ-1* KO neurons (n=20 vs 21). (**G**) Box-whisker plot quantification of the copy number of VMAT2 protein per vesicle in control and *DJ-1* KO neurons (n=558 vs 1034).

In order to obtain a more in-depth analysis of the ultrastructural organization of dopaminergic synapses, we performed a recently developed single-molecule localization microscopy (SMLM), 3D MINFLUX nanoscopy (*30, 31*) on identified individual synapses (Fig. 3B-G). Minflux requires minimal photon fluxes from the fluorophores to achieve unprecedented 3D localization precision down to ∼1 Å (angstrom). In this work, we combined Minflux with DNA PAINT, another SMLM technique, wherein the transient binding of a dye-labelled imager DNA strand on the target mimics the fluorophore blinking events that are captured in SMLM. Dopaminergic synapses were labeled using a VMAT2 monoclonal primary antibody-secondary nanobody complex carrying a docking strand, while Piccolo, a pre-synaptic scaffolding protein, was labeled using the conventional primary-secondary antibody complex. Piccolo acted as a marker to identify individual synapses and was imaged in the confocal mode prior to Minflux imaging (Fig. 3E, left). The Minflux-DNA PAINT combination drastically reduced the high background signal that are commonly associated in dense structures like synapses, and improved the localization precision in both lateral and axial dimensions, leading to a localization precision of individual fluorophores <3 nm in both axial and lateral dimensions (Fig. 3C). Overall, our Minflux-DNA PAINT imaging clearly revealed the presynaptic vesicles as clusters of fluorophores organized in 3D within presynaptic boutons (Fig. 3E, right). Such a clear 3D visualization of presynaptic vesicles allowed us to quantify parameters including total number of VMAT2-positive vesicles per synapse, vesicle diameter, and vesicular density, which are traditionally investigated using transmission electron microscopy (TEM). Furthermore, the molecular-level resolution of Minflux microscopy enabled, for the first time, detection of the copy number of VMAT2 protein per vesicle. Although no difference in the vesicle diameter (Fig. 3D, left) and the distance to the nearest neighboring synaptic vesicle (Fig. 3D, right) was observed, we found a significant reduction in the total number of VMAT2-positive vesicles per synapse (Fig. 3F), and a reduced VMAT2 protein copy number per vesicle (Fig. 3G) in *DJ-1* KO neurons over controls, indicating potential impairment of vesicular DA uptake at the *DJ-1* KO presynaptic terminals.

In summary, the insights provided by MINFLUX offer a cutting-edge approach for ultrastructural analysis of neuronal synapses, with the potential to uncover disease-related alterations at the single-molecule level. Thus, MINFLUX imaging revealed that VMAT2 is not only globally reduced at the neuronal level, as shown by whole-cell proteomics, but also at the single synapse and single vesicle level. This represents a detrimental disruption of synaptic functionality in DJ-1-deficient neurons.

### Reduced sequestration of DA revealed by false neurotransmitter assays and diminished ATP levels in DJ-1-deficient dopaminergic neurons

So far, our results uncovered diminished VMAT2 abundance at molecular, micro-, and nano-scales in *DJ-1* KO neurons when compared to the respective controls. For further experimental analysis we included another CRISPR-edited cell line and its respective isogenic counterpart by differentiating iPSCs from a PD patient carrying *PARK7/DJ-1* mutation c.192G>C (DJ-1^mut-patient^) and its CRISPR-edited control (CTRL^mut-edited^) (*32*) into midbrain dopaminergic neurons (fig. S1, S2). Using those three pairs of gene-edited DJ-1 deficient cell lines, we first confirmed the reduced abundance of VMAT2 in *DJ-1* KO neurons (Fig. 4A) as well as DJ-1-linked PD patient neurons (Fig. 4B) via immunoblotting. While VMAT2 protein levels are an important parameter for neurons that are designed to sequester (and release) DA neurotransmitter, VMAT2 transport activity is the most predictive determining factor of the efficiency of vesicular DA uptake (*33, 34*). We therefore assessed VMAT2’s activity using fluorescent false neurotransmitter in fluorometric live cell assays (*35–37*), involving the use of synthetic compounds that mimic the properties of natural neurotransmitters and are therefore valuable tools for studying neurotransmitter dynamics, vesicular release, and synaptic activity in live cells. Using FFN206 (*36*), a fluorescent false neurotransmitter probe structurally similar to DA and modified to emit fluorescence when taken up into synaptic vesicles (Fig. 4C), we found the vesicular sequestration capacity of VMAT2 to be diminished in *DJ-1* KO neurons as well as DJ-1-linked PD patient neurons (Fig. 4D), corroborating the impaired VMAT2 expression observed in these neurons. To verify that the transport is mediated through VMAT2 in this experiment, we treated control dopaminergic neurons with VMAT2 inhibitor tetrabenazine, which, as expected, led to a drastic disruption of VMAT2 activity (Fig. 4D).

**Figure 4:**
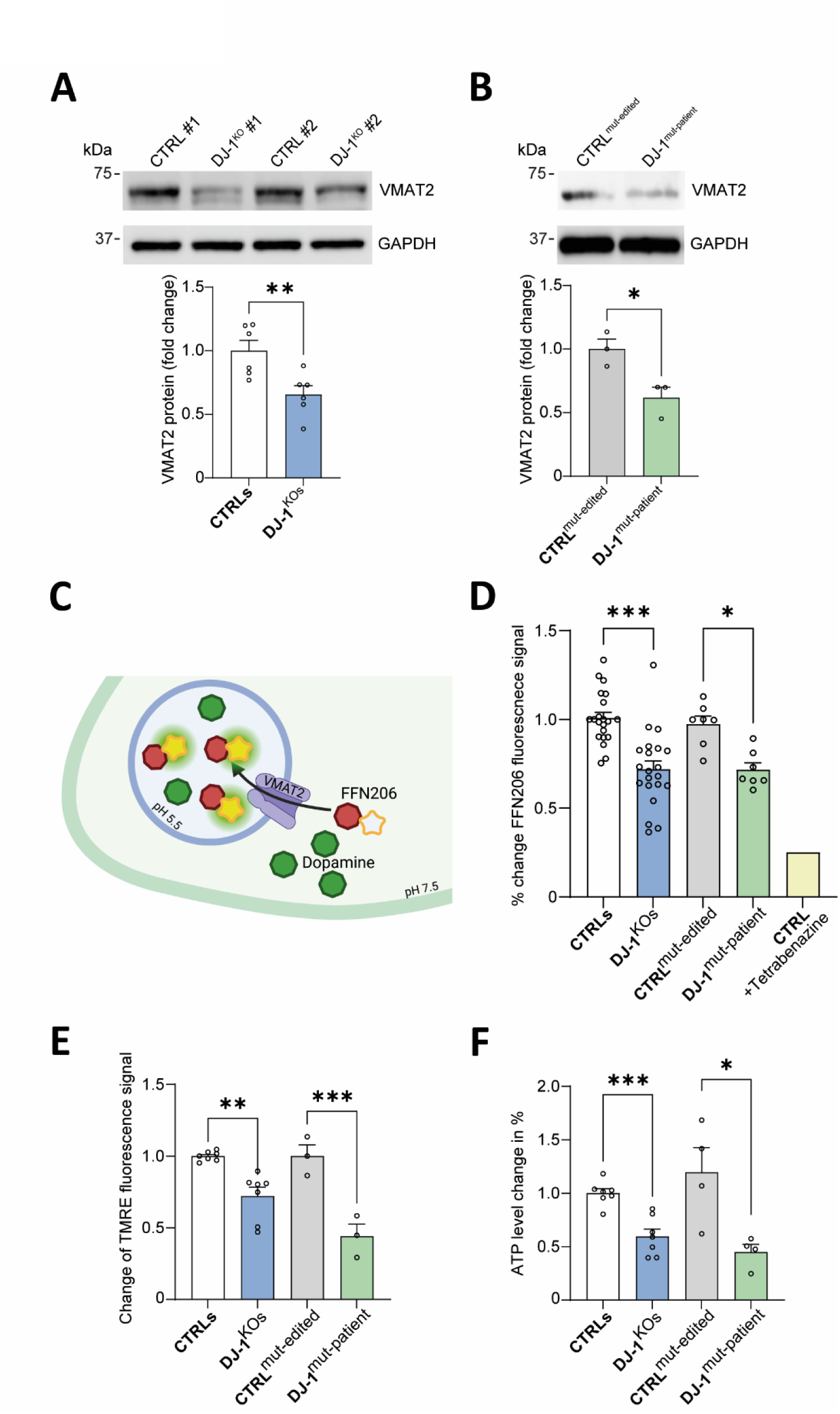
Reduced sequestration of DA due to diminished mitochondrial function and low ATP levels in DJ-1-deficient dopaminergic neurons. (**A-B**) Immunoblot analysis of VMAT2 in T-soluble neuronal lysates from (A) *DJ-1* KO and isogenic control neurons (n = 6), or (B) DJ-1-linked PD patient and isogenic control neurons (n = 3). GAPDH was used as loading control. (**C**) Schematic diagram of the false neurotransmitter illustrating the presynaptic neuron with the synaptic vesicle represented in blue, DA as green circles and the false neurotransmitter as red circles. Following vesicular sequestration of the false neurotransmitter through VMAT2, fluorescence is initiated (yellow glow attached to the false neurotransmitter red circle) through a drop in pH and can be quantified using a microplate reader. (**D**) Quantification of the dynamics of intraluminal DA loading by false neurotransmitter assay of *DJ-1* KO and isogenic control neurons, or DJ-1-linked PD patient and isogenic control neurons. Tetrabenazine was used to inhibit VMAT2 activity as control experiment (n=7-21). (**E**) Measurement of TMRE fluorescence signal of *DJ-1* KO and isogenic control neurons, or DJ-1-linked PD patient and isogenic control neurons. TMRE fluorescence correlates with mitochondrial membrane potential. The results were normalized to MitoTracker signal (n = 3-7). (**F**) Measurement of ATP levels in *DJ-1* KO and isogenic control neurons, or DJ-1-linked PD patient and isogenic control neurons (n = 4-7).

Diminished VMAT2 activity observed in the fluorometric assay could reflect either the reduced VMAT2 expression levels or an inherent impairment of VMAT2 transport activity, or both. To determine whether VMAT2 activity is, at least in part, involved in this pathology, we assessed whether its transport function is compromised in these neurons. Knowing that proper mitochondrial function and ATP supply is a prerequisite for VMAT2 functionality, we first assessed the mitochondrial health using TMRE, a mitochondrial membrane potential indicator. Quantification of the percentage of TMRE-labeled mitochondria revealed dysfunctional mitochondria with compromised membrane potential in *DJ-1* KO neurons as well as DJ-1 linked PD patient neurons (Fig. 4E). Next, we examined the levels of ATP using a fluorometric assay, and detected reduced ATP in both *DJ-1* KO and PD patient-derived neurons compared to controls, potentially reflecting the result of diminished mitochondrial function (Fig. 4F). Taken together, our results show that DJ-1-deficient midbrain neurons exhibit both a drastic deficiency of VMAT2 protein levels across different scales as well as severely impaired VMAT2 transport activity, leading to multi-level disturbance to VMAT2-dependent sequestration of DA into synaptic vesicles.

### Analysis of synaptic vesicle ultrastructure uncovers morphological abnormalities in DJ-1-deficient dopaminergic neurons

The identified numerical differences of VMAT2-positive synapses and single vesicles led us to investigate synaptic terminals at higher resolution to ultrastructurally analyze the morphology of synaptic vesicles using TEM. While control neuron terminals contained similarly sized and uniform synaptic vesicles (Fig. 5A), *DJ-1* KO neuron terminals exhibited a variety of abnormally large-sized synaptic vesicles (Fig. 5B). This observation is further supported by the Minflux data, which detected nearly twice as many abnormally sized vesicles (>100 nm in diameter) in *DJ-1* KO neurons when compared to isogenic control neurons (Fig. 5C). Interestingly, in the EM images of *DJ-1* KO neurons, we also observed the presence of intriguing tubular structures within synaptic terminals that may represent morphologically altered synaptic vesicles or tubolo-vesicular membranes resembling endosomes (*38*) (Fig. 5B), which were absent in the control neurons.

**Figure 5:**
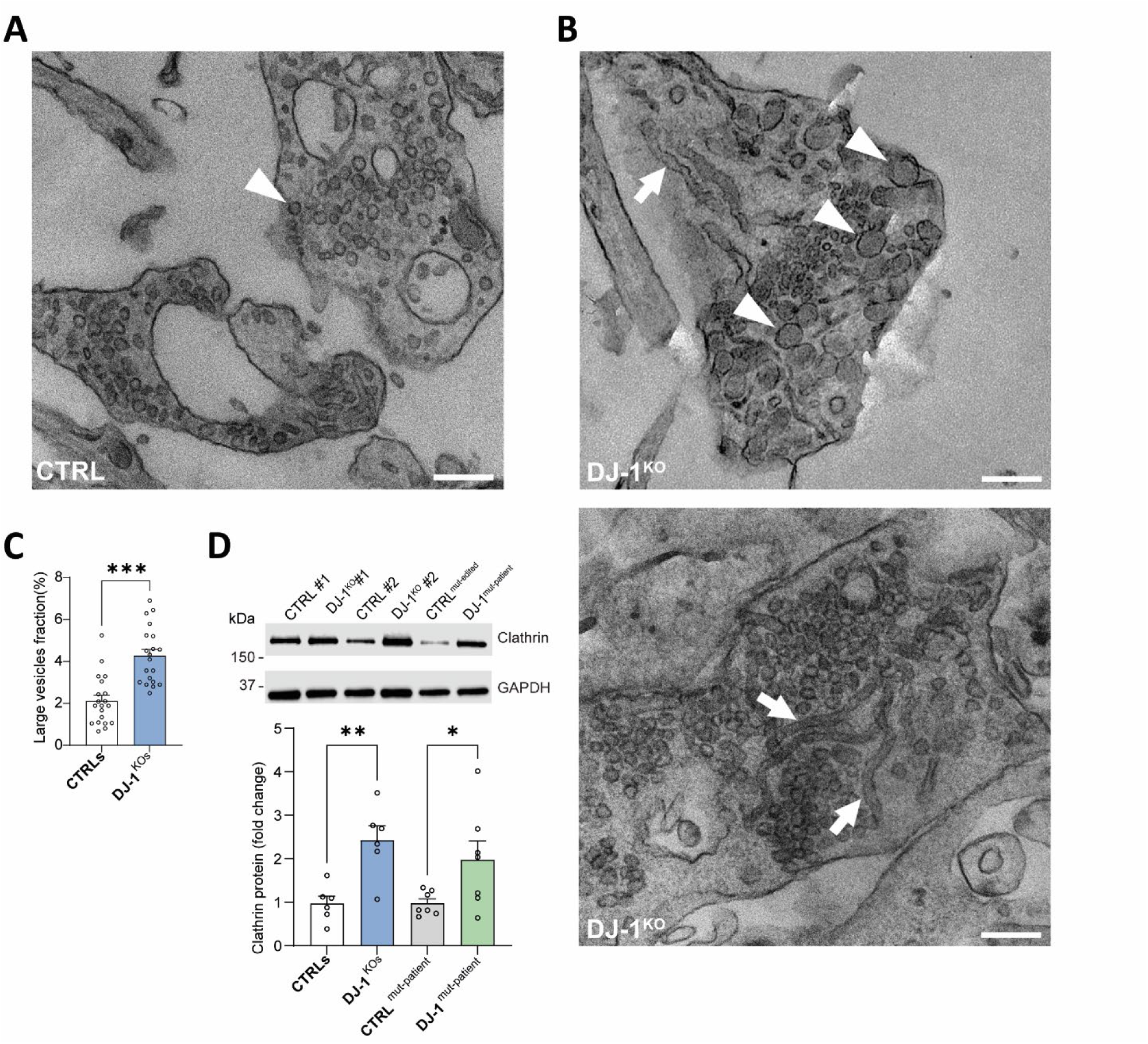
Morphological abnormalities of synaptic vesicles and excess clathrin in DJ-1-deficient dopaminergic neurons. (**A-B**) Representative transmission electron microscopy image depicting (A) regular sized synaptic vesicles (arrowhead) in control neurons, and (B) enlarged (arrowheads) or tubular (arrows) synaptic vesicles in *DJ-1* KO neurons. Scale bar, 200 nm. (**C**) Quantification of the large vesicle fraction (i.e. structures with >100 nm in diameter) in dopaminergic synapses of control and *DJ-1* KO neurons (n = 20). (**D**) Immunoblot analysis of clathrin in T-soluble neuronal lysates from *DJ-1* KO and isogenic control neurons, or DJ-1-linked PD patient and isogenic control neurons (n = 6-7). GAPDH was used as loading control.

Our results derived from EM and Minflux imaging analysis uncovered an abnormal vesicle morphology in *DJ-1* KO neuron synapses. We hypothesized that these defects originate from potential aberrant regulation in synaptic vesicle endocytosis and recycling, the pivotal local phenomenon that underlies the formation of new synaptic vesicles. In particular, the uncoating process of a newly endocytosed vesicle requires ATP-derived energy to dissociate the clathrin coat, a key protein involved in vesicle endocytosis, by retrieving vesicle components from the plasma membrane through the assembly of clathrin-coated pits (*39*), from the vesicle. Defective clathrin function can result in vesicles with abnormal shapes or sizes, disrupting their ability to efficiently store and release neurotransmitters (*40*). We assessed clathrin levels by immunoblotting and found the abundance of clathrin was approximately doubled in *DJ-1* KO and DJ-1-linked PD patient neurons compared to their respective controls (Fig. 5D). Overall, the above results suggest that, besides the deficits in VMAT2 abundance and functionality, various morphological abnormalities of synaptic vesicles were also present in DJ-1-deficient neuron synapses, possibly due to a disruption in the vesicle recycling pathways.

### Oxidized DA and α-synuclein pathology in DJ-1-deficient dopaminergic neurons

As a next step, we aimed at disentangling pathological consequences of loss of DJ-1 that may be related to the decreased abundance of VMAT2-positive vesicles and synapses, its diminished function and the observed structural vesicular abnormalities. A direct consequence of those outlined vesicular deficits is the partial loss of the capacity to sequester DA, which would in turn remain cytosolic and oxidize over time. Using a near-infrared-fluorescence (nIRF) assay to visualize oxidized DA (*41*), we found elevated levels of DA oxidation in both, *DJ-1* KO and DJ-1-linked PD patient-derived neurons (Fig. 6A). Interestingly, while the loss of DJ-1 led to a reduction of VMAT2 protein levels and DA oxidation in human neurons, lack of DJ-1 in analogous iPSC-derived mouse dopaminergic neurons generated from *DJ-1* KO mouse embryonic fibroblasts (fig. S7A to D) resulted in a similar reduction of VMAT2 protein in immunoblotting (fig. S7E), but no DA oxidation using nIRF assay (fig. S7F) compared to WT mouse iPSC-derived dopaminergic neurons. While it is known that rodents do not form NM in SN under physiological conditions (*42, 43*), and it has been previously shown that oxidized DA does not accumulate in mouse SN brain tissue or iPSC-derived mouse dopaminergic neurons(*3*), the underlying species-specific differences that lead to the lack of this human-specific pathology are still unclear. Increased levels of cytosolic DA, oxidized DA accumulation and DA-protein adduct-formation are implicated in the covalent modification and aggregation of α-synuclein, the primary structural component of Lewy bodies and a key pathological feature of PD (*44–46*). Recent studies have linked the loss of DJ-1 function to the accumulation of various forms of α-synuclein including oxidized α-synuclein (*3*), and the formation of Lewy bodies (*47, 48*). By using antibodies against all forms of α-synuclein (C20 antibody) or specific to pathogenically oxidized/nitrated (syn303 antibody) α-synuclein, we found α-synuclein to be elevated in *DJ-1* KO and DJ-1-linked PD patient neurons compared to their respective isogenic controls (Fig. 6B).

**Figure 6:**
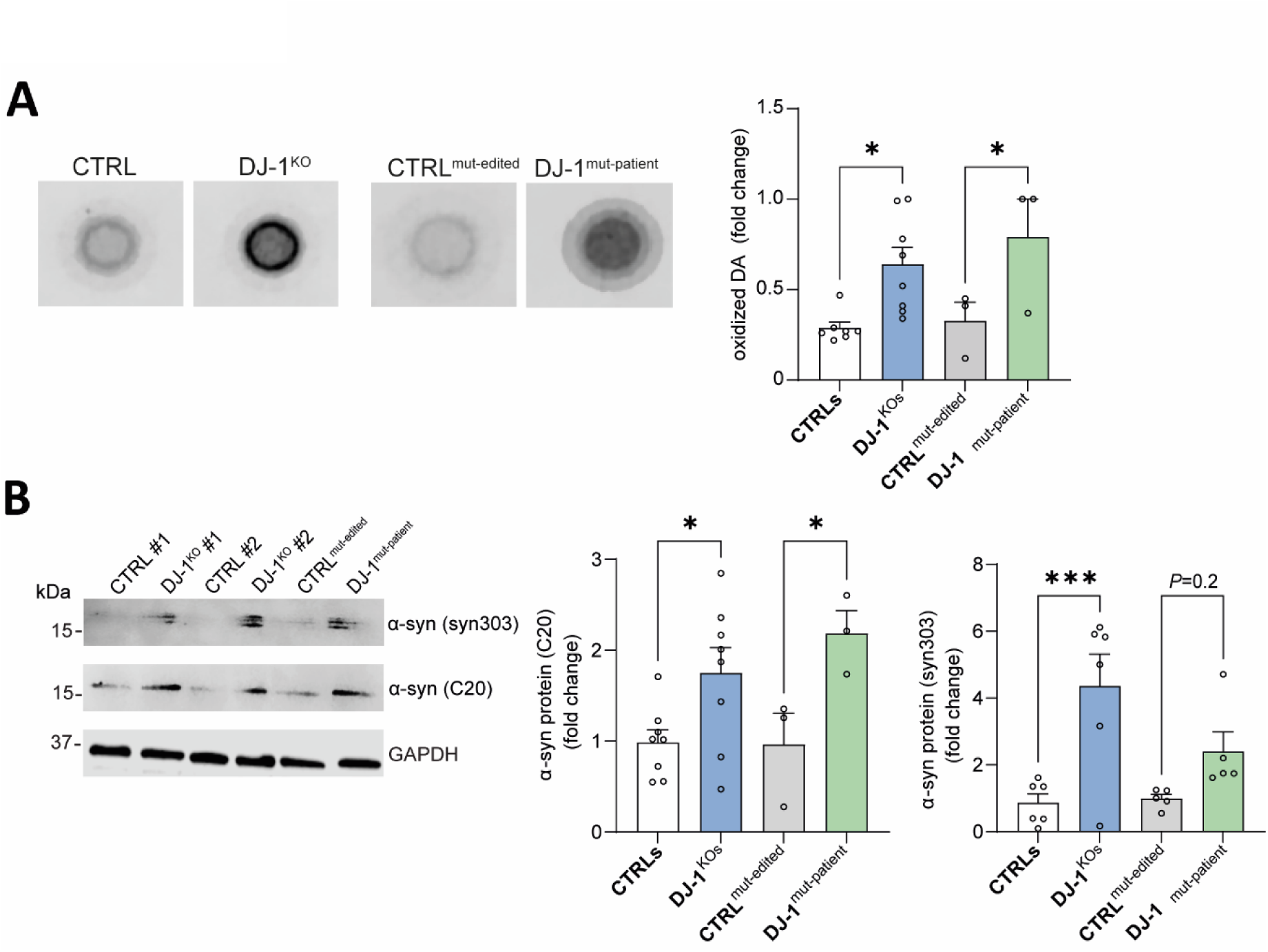
Oxidized DA and α-synuclein pathology in DJ-1-deficient dopaminergic neurons. (**A**) Representative images of oxidized DA by nIRF in *DJ-1* KO and isogenic human control neurons (left pair), or PD patient and isogenic control neurons (right pair), and quantification (n=3-8). (**B**) Immunoblot analysis of all forms of α-synuclein (C20 antibody) or oxidized/nitrated forms of α-synuclein (syn303 antibody) in T-soluble neuronal lysates from *DJ-1* KO and isogenic control neurons, or DJ-1-linked PD patient and isogenic control neurons, and quantification (n=3-8). GAPDH was used as loading control.

In sum, DA oxidation and α-synuclein pathology were found in DJ-1-deficient neurons – both phenotypes that could potentially arise from a shortage of VMAT2-positive vesicles, deficits in the functionality of VMAT2 or altered vesicle recycling. These scenarios are known to affect the proper vesicular uptake of DA from the cytosol making it prone to oxidation and potentially interfering with α-synuclein.

### ATP treatment alleviates pathologies in DJ-1-deficient dopaminergic neurons

It has been shown in various cell and animal models that the loss of DJ-1 results in mitochondrial defects, and that those deficits can potentially result in a decrease in ATP (*49–52*). Our study confirmed these discrepancies in mitochondrial function and ATP levels upon loss of DJ-1, and further highlighted consequences that are unique to human PD neurons including α-synuclein pathology and DA oxidation.

Next, as a rescue strategy, we supplemented *DJ-1* KO and DJ-1-linked PD patient neurons with ATP, directly transferred from the extracellular space into the cell by nucleoside transporters, nucleotide channels, or through micropinocytosis (*53–55*). Treatment of *DJ-1* KO and PD patient neurons with ATP led to an increase of fluorescent false neurotransmitter signal in fluorometric live cell assays indicating an increased VMAT2 transporter activity, and potential increase of DA uptake (Fig. 7A). As a proof of concept that the observed effects on replenished VMAT2 functioning were due to energy derived from supplemented ATP, we treated DJ-1-deficient neurons with ATP-γ-S (Adenosine 5’- (γ-thio)-triphosphate), a non-hydrolyzable analog of ATP, thought to fail to deliver the required energy for vesicular sequestration of DA. Indeed, DJ-1-deficient neurons supplemented with ATP-γ-S showed no change in fluorescent false neurotransmitter signal compared to untreated counterparts (Fig. 7B). We wondered whether the observed elevated levels of clathrin in DJ-1-deficient neurons would also be alleviated upon ATP supplementation. In fact, addition of ATP lowered clathrin protein levels in all DJ-1-deficient lines (Fig. 7C), possibly reflecting a restored vesicle recycling. Knowing that VMAT2 activity and vesicle uncoating of clathrin – both processes that are ATP-dependent and interrupted in DJ-1-deficient neurons – may be a potential cause for the observed DA oxidation, we treated *DJ-1* KO and DJ-1-linked PD patient neurons with ATP and measured oxidized DA by nIRF. We found that DA oxidation was successfully reduced after ATP treatment in all DJ-1-deficient lines (Fig. 7D). Importantly, ATP treatment also reduced the pathologically elevated levels of α-synuclein in all three DJ-1-deficient neuronal lines (Fig. 7E).

**Figure 7:**
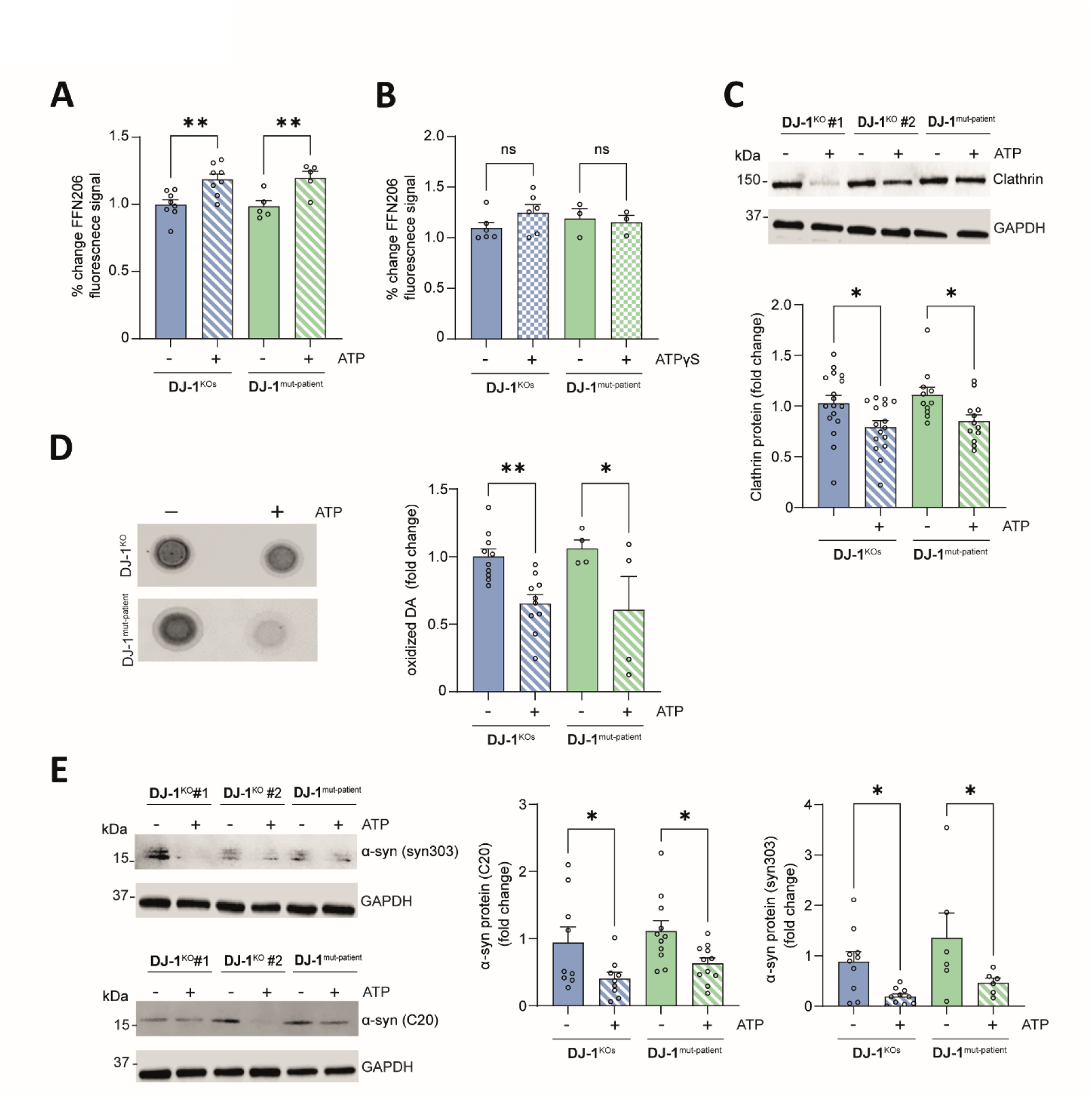
ATP supplementation lowers clathrin levels, oxidized DA and α-synuclein pathology by increasing vesicular DA sequestration in DJ-1-deficient dopaminergic neurons. (**A-B**) Quantification of the dynamics of intraluminal DA loading by false neurotransmitter assay of *DJ-1* KO and DJ-1-linked PD patient neurons treated with (A) ATP or carrier DMSO (n=5-8), or (B) ATP-γ-S (Adenosine 5’- (γ-thio)-triphosphate), a non-hydrolyzable analog of ATP, or carrier DMSO (n=3-6). (**C**) Immunoblot analysis of clathrin in T-soluble neuronal lysates from *DJ-1* KO and isogenic control neurons, or DJ-1-linked PD patient and isogenic control neurons treated with ATP or carrier DMSO, and quantification (n=12-17). GAPDH was used as loading control. (**D**) Representative images of oxidized DA by nIRF in *DJ-1* KO (top row) and DJ-1-linked PD patient neurons (bottom row) treated with ATP or carrier DMSO, and quantification (n=4-10). (**E**) Immunoblot analysis of all forms of α-synuclein (C20 antibody) or oxidized/nitrated forms of α-synuclein (syn303 antibody) in T-soluble neuronal lysates from *DJ-1* KO or DJ-1-linked PD patient neuronal lysates treated with ATP or carrier DMSO, and quantification (n = 6-11). GAPDH was used as loading control.

In summary, supplementation of ATP reduces downstream pathologies of diminished VMAT2 function, elevated clathrin and α-synuclein, as well as DA oxidation in DJ-1-deficient neurons. Overall, our results reveal that the loss of DJ-1 leads to DA oxidation and α-synuclein elevation as downstream pathologies of deficient vesicular uptake of DA that results from multiple deficits in the structure and function of VMAT2-positive synapses and vesicles, and improper uncoating of clathrin-coated vesicles in midbrain neurons (Fig. 8).

**Figure 8:**
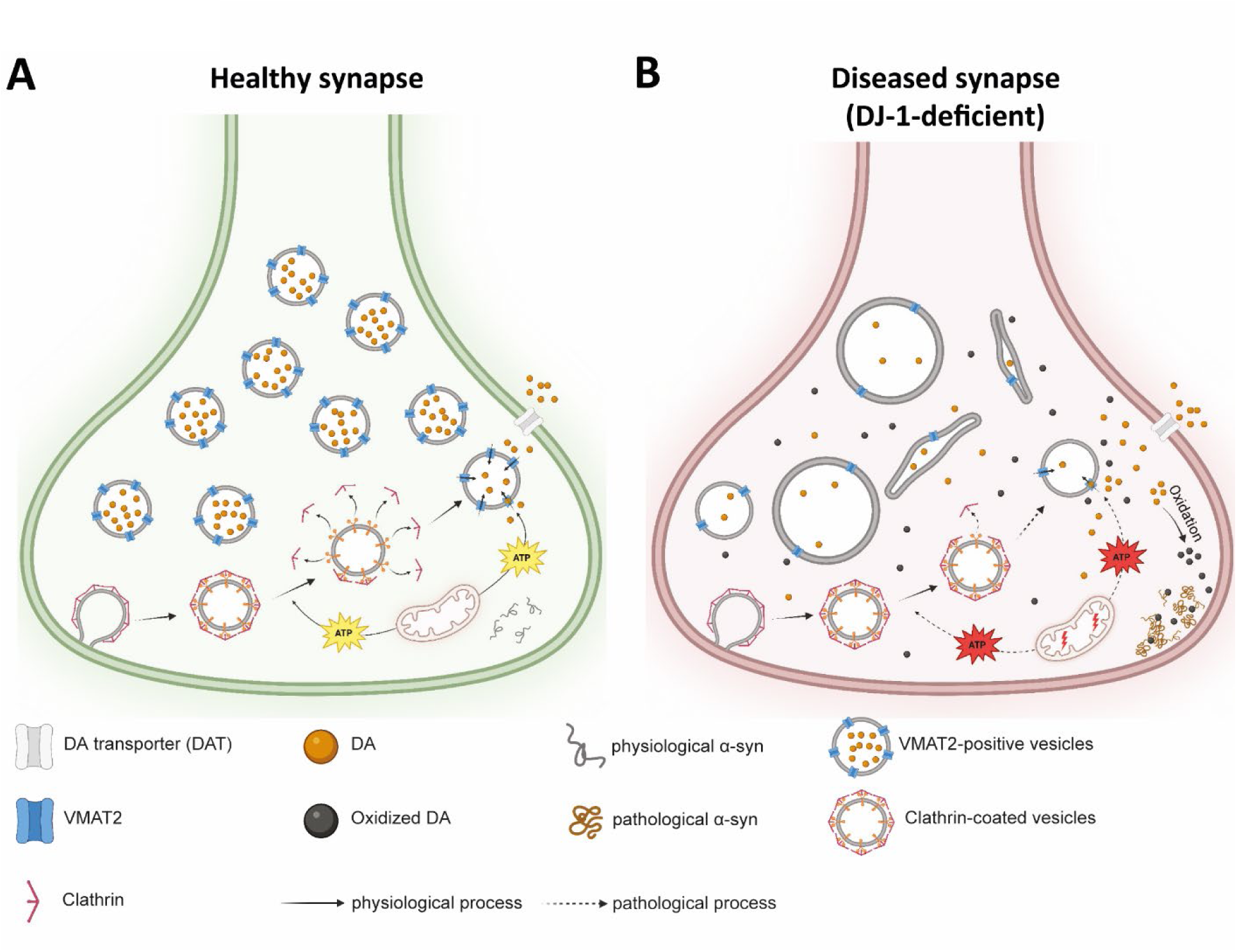
Deficient uptake of DA into synaptic vesicles of human midbrain neurons result in DA oxidation and α-synuclein pathology. (**A**) In a healthy synapse, clathrin-coated vesicles are being endocytosed, the clathrin-coat is being released and cytosolic DA is being packaged into synaptic vesicles via transport through VMAT2. Both processes, VMAT2-dependent vesicular sequestration of DA and uncoating of clathrin-coated vesicles rely on delivery of energy through sufficient ATP supply from mitochondrial sources. In this physiological situation, α-synuclein protein is present in its soluble monomeric form. (**B**) In a diseased synapse, under the pathological condition of DJ-1-deficieny, VMAT2-dependent vesicular sequestration of DA is interrupted by several synaptic pathologies. First, VMAT2 molecule number on each synaptic vesicle is diminished and VMAT2-positive vesicle number is lowered. Second, mitochondrial function is disrupted, leading to insufficient supply of ATP for processes that require energy to function. One consequence of ATP-depletion is the inadequate removal of clathrin from clathrin-coated vesicles triggering abnormally formed (enlarged and tubular) vesicle structures. The other detrimental result of lowered ATP is a functional disruption of VMTA2 activity that is incapable to transport cytosolic DA into the vesicle lumen for sequestration. These pathologies altogether lead to an increase in cytosolic DA that is rapidly converted into oxidized forms of DA, further causing neural toxicity and synaptic dysfunction, as also shown by elevated levels of pathological α-synuclein species. Conclusively, loss of DJ-1 is linked to DA oxidation and α-synuclein pathology in midbrain neurons and connects these downstream pathologies to synaptic disturbances, in particular deficits in vesicular uptake of DA and vesicle structure. Created with BioRender.com.

## Discussion

In this study, we demonstrate that disrupted vesicular sequestration of DA in DJ-1-deficient midbrain dopaminergic neurons contributes to the accumulation of oxidized DA and α-synuclein, key drivers of PD pathology. A large body of evidence links mitochondrial dysfunction to pathology (*56*), yet the precise link to DA oxidation remains unclear. Our work provides a mechanistic explanation by linking loss of ATP and vesicular dysfunction to this pathogenic process. These findings have significant implications for future therapeutic strategies targeting DA oxidation and synaptic vesicle integrity in PD.

We applied state-of-the-art imaging approaches including Minflux nanoscopy, EM, ultrasensitive DA-specific fluorescent probes, and unbiased as well as targeted proteomic analyses, to quantify disturbances of synaptic vesicle function upon loss of the PD-associated protein DJ-1 in three pairs of CRISPR-edited cell lines with their respective isogenic counterparts. Whole-cell proteomics and MS following BN-PAGE and IP all indicated synaptic vesicle integrity to be affected in neurons lacking DJ-1 with VMAT2 being a key protein found to be substantially reduced.

The role of VMAT2 in DA regulation and its impact on neurodegeneration has been widely debated (*57*). Notably, VMAT2 has been shown to be markedly reduced in idiopathic PD patient brain (*58*), and regions more resilient to degeneration in PD such as the neighboring ventral tegmental area (VTA) tend to exhibit higher VMAT2 expression levels suggesting a neuroprotective role (*59*). Importantly, the latter study emphasizes an inverse correlation between the accumulation of the characteristic pigment NM – which forms when excess cytosolic DA is present (*10*) – and VMAT2 levels suggesting that human nigral neurons accumulate the most NM pigment, in part, because they have the least VMAT2 protein. In line with this, a specific VMAT2 gain-of-function haplotype has been associated with a reduced risk for the development of PD (*60*).

It has been shown that siRNA-mediated reduction of DJ-1 levels resulted in reduced VMAT2 levels, though these results were solely collected under knockdown conditions in a human neuroblastoma cell line under naïve, undifferentiated conditions (*61, 62*). Nevertheless, these results suggest that, at least in neuroblastoma cells, DJ-1 positively regulates VMAT2 expression and suggests a more general role of DJ-1 in regulating VMAT2 levels. This rather universal association between DJ-1 and VMAT2 may also explain our results on comparable VMAT2 reduction in human and mouse iPSC-derived *DJ-1* KO neurons. However, interestingly, while VMAT2 deficiency was similar between the species, only the loss of DJ-1 in human, but not mouse, dopaminergic neurons resulted in an accumulation of oxidized DA. This is in line with previous studies (*3*) and underscores the notion that also major PD pathologies, such as significant neuron loss, are absent in *DJ-1* KO mouse models (*63–65*). One potential reason for DA oxidation reflecting a species-specific difference between human and mouse midbrain dopaminergic neurons may be the ability of the murine system to compensate for a reduction in VMAT2, through other, yet unknown, mechanisms. While more research needs to be done to understand the association between DJ-1 and VMAT2 in disease-relevant model systems, our own findings strengthen the link between DJ-1 deficiency, diminished VMAT2-dependent vesicular sequestration, and toxic DA oxidation in synapses of vulnerable human midbrain neurons.

Besides the observed VMAT2 transporter (activity) disturbances, another intriguing phenotype of synaptic integrity was the disrupted vesicle morphology. While the underlying reasons for the observed irregular vesicle formations are unclear and need more investigation, the overabundance of clathrin in DJ-1-deficient neurons suggest the inadequate removal of clathrin from clathrin-coated vesicles triggering abnormally formed (enlarged and tubular) vesicle structures (Fig. 8). Clearly, both pathologies – VMAT2 deficiency and irregular vesicle morphology – impair the uptake of DA into vesicles of DJ-1-deficient midbrain synapses leading to the buildup of unsequestered DA in the cytosol, which is susceptible to oxidation.

Oxidized DA is a highly neurotoxic compound and has been shown to be able to modify proteins, including α-synuclein (*44, 45*). In fact, in our study we also found forms of α-synuclein, including the oxidatively modified forms, to be increased in DJ-1-deficent neurons. α-Synuclein, a natively unfolded protein predominantly localized to the presynaptic terminal, has been extensively studied in PD (*66*). Under pathological conditions, α-synuclein is often found abnormally phosphorylated, ubiquitinated, oxidized and nitrated (*67*), and these toxic α-synuclein species impair cellular function and survival (*68*). There is evidence for a role of DJ-1 in inhibiting the initial aggregation of α-synuclein monomers through its molecular chaperone-like activity (*69, 70*). Furthermore, research has shown that loss of DJ-1 leads to an increase in aggregated α-synuclein levels in both cell and animal models of PD (*71*). In this regard, it is of interest to mention that the presence of pathologic α-synuclein oligomers has been shown to initiate downregulation of synapsins, i.e. SYN1 and SYN2, primarily involved in the regulation of neurotransmitter release and synaptic vesicle dynamics (*72*). Interestingly, our proteomic whole cell analysis revealed both SYN1 and SYN2 to be diminished in DJ-1-deficient neurons. This highlights the possibility of a toxic feedforward loop of deficient vesicular sequestration leading to DA oxidation and α-synuclein accumulation, that in turn may affect vesicular uptake of DA by affecting synapsin levels, and, over time, enhance the level of neuron vulnerability. In addition, proteomics revealed reduced calcium ion binding in *DJ-1* KO neurons, suggesting impaired calcium handling. This aligns with the findings of Guzman et al. (2010) (*13*), who demonstrated that DJ-1 attenuates oxidant stress generated by calcium-dependent pacemaking activity in dopaminergic neurons. Our data may indicate that loss of DJ-1 compromises calcium buffering or signaling, possibly making neurons more vulnerable to oxidative stress.

The question remained how such a neurotoxic feedforward loop was initiated in DJ-1-deficient neurons. There is broad evidence in the literature for disturbed mitochondrial dysfunction concomitant with reduced ATP levels upon loss of DJ-1 in various experimental models (*13, 49–52*). However, this loss of ATP is specifically problematic in dopaminergic neurons that rely on this source of energy for additional resources, including VMAT2 activity and uncoating of clathrin-coated vesicles. Indeed, all DJ-1 mutant lines in our study presented with a reduced mitochondrial membrane potential and ATP. Therefore, as a proof-of-concept experimental model, we aimed at interrupting the initiation of this toxic cycle and supplemented DJ-1-deficient neurons with ATP. This strategy uses the remaining, well-functioning VMAT2 molecules, to package cytosolic DA into synaptic vesicles reducing the risk of oxidation. Indeed, ATP treatment successfully rescued the observed pathologies, including diminished VMAT2 transporter activity, DA oxidation and elevated α-synuclein levels.

Mitochondrial dysfunction has been described as one of the key pathological drivers of PD, in both familial and sporadic forms (*73*). Importantly, compounds, such as MPTP, rotenone and paraquat, that are known to interfere with mitochondrial function and induce PD-like symptoms have been used to generate neurotoxic models of PD since decades (*56*). Notably, energy defects and diminished ATP production have been observed in PD patients and linked to α-synuclein aggregation (*74, 75*). In fact, mitochondrial dysfunction has been detected even prior to α-synuclein aggregation or neurodegeneration, suggesting it being a potential causal trigger or even an initiator of the disease pathology (*76*), and of the toxic cascade described in our study (Fig. 8). While supplementation of ATP has been used in our study as an experimental model, the improvement of mitochondrial function has been examined as a strategy for reducing PD pathology and benefiting neuron survival. However, most clinical trials targeting brain energetics were unable to meet key clinical endpoints (*56*). Still, much of the research is in early stages, and further studies are needed to fully understand how mitochondrial modulators can be effectively applied in clinical settings.

Patient-derived neuronal models are essential for exploring the disease mechanisms underlying neuron pathophysiology and evaluating potential therapeutic strategies. To gain a more comprehensive understanding of neuronal dysfunction, advanced human culture systems are needed to better replicate the intricate cellular architecture of the brain. Our study marks a significant step in unravelling the complexities of DA metabolism in human midbrain neurons and open new avenues for the development of interventions tailored to dysfunctional pathways of vesicular DA sequestration.

## Materials and Methods

### Culture and characterization of human iPSCs, and differentiation into midbrain dopaminergic neurons

We used iPSC lines that were already generated from fibroblasts from one DJ-1-linked familial PD patient (DJ-1^mut-patient^) carrying the homozygous c.192G>C mutation in *PARK7* (DJ-1) that was clinically characterized previously (32) and its isogenic CRISPR-edited control (CTRL^mut-edited^), and from two separate healthy control subjects (CTRLs) (3, 77–79) and their isogenic PARK7/DJ-1 KO counterparts (*DJ-1* KOs) (3). CRISPR-Cas9 gene editing strategies were described previously (3, 32). The study was approved by the Ethics Committee of the Medical Faculty of LMU Munich, Germany (approval number 25-0153). All patients and controls gave written and informed consent.

iPSC cultures were kept on plates coated with Cultrex (R&D Systems, #3434-005-02) and the antibiotic-free mTESR^TM^ Plus medium (Stem Cell Technologies, #100-0276) was replenished every two days. Cells were passaged manually every 5-7 days. Each line was assessed for expression of pluripotency markers by immunocytochemical analysis (Oct4 (Abcam, #ab19857, 1:200), SOX2 (Abcam, #ab79351, 1:200), Nanog (Abcam, #ab80892, 1:200), SSEA-4 (Millipore, #MAB4304 at 1:200), as well as for in vitro differentiation via embryoid body formation (see below). Additionally, all lines were regularly screened for mycoplasma contamination.

The terminal differentiation of human iPSCs into midbrain dopaminergic neurons was conducted according to previously established protocols (3, 80). In summary, once the cells achieved over 95% confluence (Day 0 = d0), a differentiation medium comprising Knockout DMEM medium (Thermo, #10829-018), KnockOut™ Serum Replacement (KOSR) (Thermo, #10828-028), 1% L-Glutamine, 1% PenStrep, 1% MEM Non-Essential Aminoacids (Thermo, #11140-050) and 1mM beta-mercaptoethanol (KSR media) was introduced. From days 5 to 10, the cells were transitioned to Neurobasal media (Thermo, #21103049) enriched with NeurocultTM SM1 (Stem Cell Technologies, #5711), 1X penicillin-streptomycin, and 1X L-glutamine (NbSm1 media). During days 11 to 14 (d11-d14), the cells were passaged en bloc in 1-2mm pieces and transferred onto 10cm dishes coated with poly-d-lysine (PDL)/laminin, continuing to be maintained in NbSm1 medium. Between days 25 and 30 these chunks were harvested with accutase and passaged onto PDL/laminin-coated culture dishes for the final experiments. Neuralization factors were freshly added from days 1 to 40-50, after which the neurons were kept in NbSm1 medium.

The identification of dopaminergic lineage neurons was achieved through immunocytochemical staining for DA, midbrain, and neuronal markers at day 30 of differentiation. Only fully mature midbrain dopaminergic neurons at day 70 of differentiation were utilized for experimental analysis.

### Embryoid body formation

Human iPSC colonies were rendered as single cell suspension by using Accutase (Sigma). Cells were centrifuged (300g x4 mins), resuspended and seeded into 96 wells ultra-low attachment spheroid microplate (Corning, #4520) in E6 medium at a density of 1.5x10^4^ cells/well. Immediately after seeding the cells, the plates were briefly centrifuged (100g x3 mins) to allow them to evenly spread at the bottom of the wells. On day 7, formed embryoid bodies (EBs) were plated onto PDL/laminin-coated plates in KO DMEM F12 medium (Thermo Fisher Scientific) supplemented with 10% heat-inactivated FBS (inactivation performed at 56° for 30 min). After additional 7 days of spontaneous differentiation, attached EBs were harvested and qRT-PCR analysis was performed for markers of the three germ layers.

### RNA extraction and qRT-PCR

qPCR was performed as previously described (*77*). Briefly, Total RNA was extracted and isolated using the RNeasy Mini Kit (Qiagen) following vendor’s protocol, and RNA concentration was assessed using a NanoDrop 2000C (Thermo Scerntific). Complementary DNA (cDNA) was synthesized with 500 ng of RNA following vendor’s protocol (iScript cDNA Synthesis Kit Bio-Rad). SYBR green qPCR was conducted using the Fast SYBR™ Green Master Mix (Applied Biosystems) following standard protocol and ran in triplicate on the StepOnePlus Real-Time PCR System (Applied Biosystems). Data were quantified using the ΔCt-method and normalized to GAPDH as a reference gene.

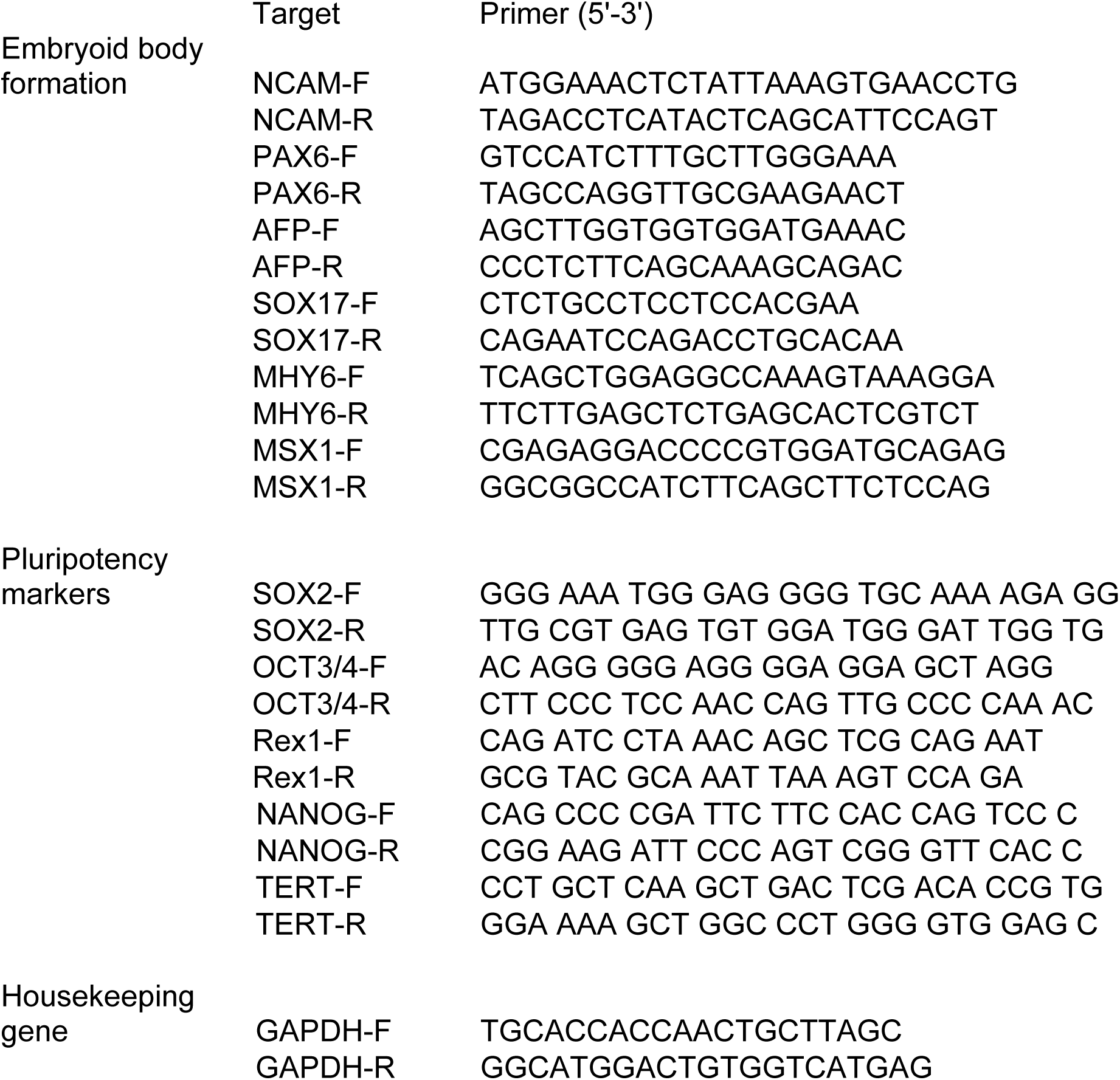

### Reprogramming of mouse embryonic fibroblasts to iPSCs

Mouse embryonic fibroblasts (MEFs) were received from *DJ-1* KO and WT mice described earlier (*78*). MEFs were reprogrammed into iPSCs using the CytoTune-iPS 2.0 Sendai reprogramming kit (Thermo Fisher, #A16518) following standardized vendor’s protocol. iPSCs generated from MEFs derived from three WT mice were expanded and one clone per animal used for experiments. iPSCs generated from MEFs derived from one DJ-1 KO mouse were expanded and two clones of this iPSC lines used for experiments.

### Mouse iPSC culture and neural differentiation

Mouse iPSCs were cultured as floating sphere on low attachments plates located in an incubator provided with a constant shaking plate. iPSCs were cultured in iPSCs media (Knockout DMEM F12, 10% KOSR, 1% L-Glutamine, 1% PenStrep, 1% MEM Non-Essential Aminoacids, 55 uM beta-mercaptoethanol) supplemented with ESGRO 2i-LIF (Millipore, #ESG1121, 1:1000). Media was changed daily and cells were passaged every 3 to 4 days as single cells.

Mouse neuronal differentiation was performed following published protocols (*79*) with some modifications. Briefly, iPSCs ready for passage were rendered into single cell suspension using TrypLE (Thermo Fisher Scientific). 1-1.5x10^6^ iPSCs were seeded onto low attachment 10 cm dishes in iPSCs media without ESGRO 2i-LIF, and positioned in an incubator provided with a constant shaking plate for 3 days to form embryoid bodies (EBs). On day 4, EBs were gently collected and seeded onto 10 cm dishes coated with PDL-laminin in iPSCs media without ESGRO 2i-LIF. The following day, once EBs were attached to the plate, media was changed to differentiation media (DMEM F-12, 1% N2 supplement, 1% PenStrep, 1% L-Glutamine and 1 ug/ml laminin) supplemented with 10 ng/ml bFGF2 (R&D, #100-18b), 100 ng/ml FGF8b (R&D, #423-F8-025/CF) and 10 nM SAG 1.3 (Sigma-Aldrich, #566660). Four days later, EBs were dissociated into single cells using TrypLE and seeded onto PDL-laminin coated plates. After two days, bFGF2, FGF8 and SAG 1.3 were removed from the media, and 200 µM of ascorbic acid (Sigma-Aldrich, #A5960) was added. After six days, media was further supplemented with 10 ng/ml BDNF (R&D, #248-BDB-250/CF) and 10 ng/ml of GDNF (R&D, #212-GD-01M) and cells were maintained until fully differentiated (= 30 days of differentiation).

### Sample Preparation for LC-MS/MS

A modified procedure for single-pot solid-phase enhanced sample preparation (SP3) was used (*80*). In short, 15 µg of protein lysate was mixed with *MgCl_2_* to a final concentration of 10 mM. After adding 25 U Benzonase (Sigma-Aldrich) samples were incubated for 30 minutes at 37°C. Proteins were reduced and alkylated by adding dithiothreitol (Biozol) to a final concentration of 10 mM, and then incubated for 30 minutes at 37°C. Iodoacetamide (Sigma-Aldrich) was then added to a final concentration of 40 mM, and the reaction was quenched by adding another portion of dithiothreitol. Proteins were then bound to 40 µg of a 1:1 mixture of hydrophilic and hydrophobic magnetic Sera-Mag SpeedBeads (Cytiva) with a final concentration of 80% ethanol (Sigma-Aldrich) for 30 minutes at room temperature. 200 µL of 80% ethanol was used to wash the beads four times using a Dynamag-2 magnetic rack (Thermo Fisher Scientific). 190 ng LysC and 190 ng trypsin (Promega) were added to 40 µL of 50 mM ammonium bicarbonate for proteolytic digestion, and the mixture was then incubated for 16 hours at room temperature. The supernatants were dried by vacuum centrifugation after being filtered using 0.22 µm spin-filters (Costar Spin-X, Corning). Twenty microliters of 0.1% formic acid were used to dissolve the dehydrated peptides. The Qubit protein assay (Thermo Fisher Scientific) was used to quantify the peptide content following proteolytic processing.

### LC-MS/MS analysis

The LC-MS/MS proteomic analyses were conducted on a nanoElute system (Bruker Daltonics) that was online coupled with a timsTOF pro mass spectrometer (Bruker Daltonics) equipped with a column oven. 350 ng of peptides were separated on a 15 cm (75 µm ID) column self-packed with ReproSil-Pur 120 C18-AQ resin (1.9 µm, Dr. Maisch GmbH) using a 90-minute binary gradient of water and acetonitrile supplemented with 0.1% formic acid at a flow rate of 300 nL/min and a column temperature of 50°C.

Data independent acquisition Parallel Accumulation Serial Fragmentation (DIA-PASEF) was used. For peptide fragment ion spectra, 34 consecutive DIA windows with a width of 26 m/z were placed after one MS1 complete scan. These windows overlapped by 1 m/z, spanning a scan range of 350 to 1200 m/z. Each ramp had two windows examined, and the ramp time was set at 100 ms. This resulted in a total cycle time of 1.9 s.

### Data analysis of LC-MS/MS results

Label-free quantitation (LFQ) of proteins was carried out using the DIA-NN (v 1.8) (*81*). A library-free search was conducted against a database of common contaminants from Maxquant (246 entries) and a canonical one protein per gene reference sequence database of humans that was downloaded from UniProt (download date: 2023-03-01, 20603 entries). Trypsin was classified as a protease, and two missed cleavages were permitted. Carbamidomethylation of cysteines was classified as a permanent modification, while oxidation of methionines and acetylation of protein N-termini were classified as variable modifications. For the identification of proteins and peptides, a 1% FDR threshold was used.

The program Perseus (v 1.6.2.5) (*82*) was used to analyze the output. For protein LFQ, two identified peptides were necessary. For statistical testing, the protein LFQ intensities had to be log2 converted and have at least three valid values per experimental group. A Student’s t-test was used to compare protein abundance changes between the two experimental groups based on log2 LFQ intensity. Differentially expressed proteins were defined by a p-value <0.05 and a protein log2 fold change larger than 0.5 or smaller than -0.5 for both *DJ-1* KO lines, respectively.

Gene Ontology (GO) analysis was performed using the web-based program DAVID version 6.8 (*83, 84*). Gene lists for either up- or downregulated proteins were compared to all proteins detected in LC-MS/MS in all cell lines. Using EASE Score of 0.05, otherwise the default parameters, functional annotation charts were generated for molecular functions (GOTERM_MF_DIRECT), biological processes (GOTERM_BP_DIRECT), and cellular components (GOTERM_CC_DIRECT).

### Blue-Native Polyacrylamide Gel Electrophoresis (BN-PAGE)

BN-PAGE was performed largely according to the manufacturer’s instructions and, unless noted, BN-PAGE reagents were obtained from Invitrogen. Snap-frozen cell pellets from control and *DJ-1* KO dopaminergic neurons were resuspended and lysed in cold BN-PAGE lysis buffer (50 mM Tris, 1% (w/v) digitonin, pH 7.40, freshly supplemented with Protease and Phosphatase Inhibitor Cocktail (PPhI), Sigma-Aldrich, PPC1010, 100x), incubated 30 mins on ice, then spun at maximum speed in a tabletop centrifuge (15 mins). Total protein concentration in the lysate supernatants was quantified via BCA (BC Assay protein quantitation kit, Interchim) and the lysate was aliquoted in single-use aliquots and snap-frozen in liquid nitrogen. On the day of use the lysate aliquots were supplemented with NativePAGE Sample Buffer (4X) and NativePAGE 5% G-250 Sample Additive (to 0.25%), spun in a tabletop centrifuge to remove aggregates, and loaded on a 4-16% NativePAGE Bis-Tris Gel. The gel was run at 4°C at 150 V (60 mins) and then at 250 V (90 mins), and, when Western blotting, the cathode buffer was switched from dark blue (0.02% Coomassie G-250) to light blue (0.002% Coomassie G-250) when the sample dye front reached about 1/3 of the gel length. For blotting, BN-PAGE transfer buffer (25 mM bicine, 25 mM Bis-Tris, 1 mM EDTA) was used as a buffer system and the gel was transferred on 0.45-µm PVDF membranes (Immobilon-P, Sigma-Aldrich) at 4°C (25 V, 90-120 mins). After blotting, the membrane was fixed in 8% acetic acid (15 mins), washed extensively with methanol, then dried and rehydrated in TBS-T. From this point on, Western blotting followed the SDS-PAGE WB protocol, with chemiluminescent detection.

### MS sample preparation and data collection

Protein complexes from control and *DJ-1* KO neurons were separated by BN-PAGE as described in methods for “Blue-Native Polyacrylamide Gel Electrophoresis (BN-PAGE)“. Following electrophoresis, marker lanes were visualized with Coomassie blue stain and bands of interest were excised using sterile scalpels. The bands were chopped in smaller pieces and transferred onto a 96 well plate. Gel pieces were washed with 50 mM ammonium bicarbonate (ABC)/50% ethanol buffer, followed by dehydration with absolute ethanol, reduction of proteins with 10 mM Dithiothreitol (DTT) in 50 mM ABC at 56°C for 45 min and alkylation of proteins with 55 mM iodoacetamide in 50 mM ABC at room temperature for 30 min. Gel pieces were once again washed and dehydrated (twice for 15 min) as above followed by overnight trypsin digestion (12 ng/µl trypsin in 50 mM ABC, and 40 μl trypsin solution/well) at 37°C. Peptides were step-wise extracted from gel pieces with 30% acetonitrile/3% trifluoroacetic acid (TFA) and 70% acetonitrile and 100% acetonitrile. Combined supernatants were dried down, mixed 1:1 with 5% acetonitrile/1% TFA and loaded on pre-conditioned custom-made stage tips (2× C18 disks). Stage tips were washed with the 0.1% Formic acid and peptides eluted with 80% acetonitrile/ 0.1% formic acid. Peptide mixtures were separated using an Easy-nLC1200 liquid chromatograph (Thermo Scientific) followed by analysis on a Q Exactive HF mass spectrometer (Thermo Scientific). MS data was processed with MaxQuant (version 2.6.7.0) and analyzed with Perseus (version 2.1.3.0). Matches to common contaminants, reversed identifications, and identifications relying solely on site-specific modifications were excluded before further analysis. After log2-transformation of Label-Free Quantification (LFQ) intensities, proteins were filtered to include only those detected in at least three out of four replicates in at least one group (control and *DJ-1* KO). Two-sample student’s t-test was performed comparing control vs *DJ-1* KO to identify significantly changed proteins.

### Immunoprecipitation (IP)

For the immunoprecipitations, snap-frozen dopaminergic neuron pellets were resuspended and lysed in cold IP lysis buffer (50 mM Tris, 150 mM NaCl, 1% NP-40, 0.15% (w/v) bovine serum albumin, 10% (w/v) glycerol, 100x PPhI, pH 8.00), incubated 30 mins on ice, then spun at maximum speed in a tabletop centrifuge (15 mins). Pooled lysate supernatants of three 6-well-plate wells were used as the input for each IP reaction and diluted to 1 mL in IP wash buffer (10 mM Tris, 140mM NaCl, 0.1% n-Octyl-β-D-glucopyranoside, 5 mM EDTA, 100x PPhI, pH 8.00). 5 µg of anti-PARK7/DJ1 antibody (EP2815Y, abcam) were then added to each tube and the IP reactions were rotated overnight at 4°C. On the next day, using wide-bore tips, 40 µL (80 µL of slurry) of Pierce Protein A Agarose beads (Thermo Scientific) were added to each reaction and the tubes were rotated for two more hours at 4°C. The beads were then spun in a cooled tabletop centrifuge (2500 rcf, 5 mins), and, after removal of the immunodepleted supernatant, washed with IP wash buffer. Beads were washed 4x with IP wash buffer and 2x with 50 mM ammonium bicarbonate buffer, pH 8.00, then dried, snap-frozen in liquid nitrogen, and stored at -80°C. On-bead digestion and mass spectrometry were performed as previously described (https://pubmed.ncbi.nlm.nih.gov/32182352/).

### Immunofluorescent staining and image analysis

Immunocytochemistry was performed using an adapted version of previously published protocol (*77*). Cells were fixed in 4% Paraformaldehyde for 15 minutes, washed three times in PBS and then permeabilized for 30 minutes with 0.3% Triton-X100. Following permeabilization, coverslips were blocked with 0.3% Triton X-100 in 5% normal goat serum and 1% BSA in PBS for 1 hour. Coverslips were incubated overnight with the respective primary antibody against SOX2 (Abcam, #AB79351; 1:100), Nanog (Abcam, #AB80892; 1:100), OCT4 (Abcam, #AB19857; 1:100), SSEA4 (Merck, MAB4304; 1:100), TuJ (ß-III-Tubulin) (Biolegend, #801201; 1:100; Biolegend, #802001; 1:100), LMX1a (Abcam, #ab10533; 1:100), FOXA2 (Santa Cruz, #sc-101060; 1:100), TH (Tyrosine Hydroxylase) (Merck, #A657012; 1:100). The day after coverslips were washed three times with PBS, and incubated for 1 hour at room temp with the respective secondary antibodies against anti-mouse or anti-rabbit Alexa Fluor 488 or 568 (Invitrogen, #A11034; 1:1000; Invitrogen, #A11031; 1:1000). Secondary antibody was washed three times with PBS and coverslips were incubated with DAPI for 5 minutes. Coverslips were then washed three times in PBS and once in sterile H2O before being mounted on ProLong mounting media (Thermo).

For mouse iPSCs cells, spheres were collected at the moment of passage, centrifuged at 150 g x 2 minutes, resuspended and fixed in 4% PFA for 30 minutes with gentle shaking. After fixation, spheres were washed 3x in PBS (centrifugation – resuspension), and then permeabilized for 30 minutes with 0.3% Triton-X100 in gentle shaking. After permeabilization, spheres were blocked with 0.3% Triton X-100 in 5% normal goat serum and 1% BSA (1:1 ratio) in PBS for 1 hour in gentle shaking. Spheres were then incubated overnight in gentle shaking with the respective primary antibody against SOX2 (#AB79351 at 1:100), Nanog (Abcam #AB80892 at 1:100), OCT4 (Abcam #AB19857 at 1:100), SSEA1 (Merck, MAB4301; 1:100). The day after sphere were washed three times with PBS, and incubated for 1 hour at room temperature with the respective secondary antibody against anti-mouse or anti-rabbit Alexa Fluor 488 or 568 (Invitrogen, #A11034; 1:1000; Invitrogen, #A11031; 1:1000). Spheres were washed three times with PBS and incubated with DAPI for 5 minutes at room temperature in gentle shaking. Spheres were then washed three times in PBS and once in sterile H2O before being placed as drops on glass microscope glasses. Microscope glasses were quickly placed on a heated thermo block until water evaporated, and then a glass coverslip was mounted using the ProLong mounting media (Thermo).

Images were captured using confocal microscope (Leica Stellaris 5, Leica Microsystems, Wetzlar, Germany) provided with a dry 20x objective, or an oil immersion 40x and 63x objective. For each coverslip at least three fields of view were examined.

### Labeling of neurons for confocal and Minflux-DNA PAINT imaging

iPSC-derived midbrain dopaminergic neurons from both *DJ-1* KO and their CRISPR-edited control lines were grown on 24-mm coverslips. Prior to labelling, the neurons were fixed at day 70 of differentiation using 4% PFA for 15 min and the coverslips were washed twice, permeabilized for 5 min (0.3% TritonX-100; Thermo Fisher Scientific, A16046-AE), blocked with 5% normal goat serum for 1 h and incubated with primary antibodies for VMAT2 (Abcam, #ab259970), VGLUT (SySy, #135408), VGAT (SySy, #131008) and piccolo (Synaptic Systems, #142104) overnight. Next day, for confocal imaging, primary antibodies were removed, washed thrice, and incubated with dye-labeled secondary antibodies (Abcam, #ab150079, #ab150185) for 1 h. Afterwards, the coverslips were washed thrice and the samples were imaged under confocal microscopy within 1 day. For Minflux imaging, after the primary antibody incubation, the neurons were washed and incubated with nanobodies coupled to a specific single-stranded DNA docking strand (Massive Photonics, MASSIVE-SDAB2-PLEX) for 1 h at room temperature. Finally, the samples were washed thrice and post-fixed using PFA, followed by a washing step using PBS.

### Minflux imaging

Before imaging, each coverslip was incubated with fiducial gold beads (150 nm; BBI solutions, EM.GC20/7) for 5–10 min and the non-immobilized beads were removed by washing with PBS. The fiducial beads ensured nanometer scale stability of the sample during measurements by an active stabilization system that detected the 3D positions of the beads by a separated beamline and coupled with a galvanometer for real-time drift correction. Next, single stranded DNA imager strands that are complimentary to the docking strands (Massive Photonics) was added to the sample and transferred to the microscope. Initially, the neurons were imaged in confocal mode and individual synapses were identified using the punctate piccolo signal, which were chosen as specific regions of interest (ROIs) (<2.25 µm^2^ area). Each ROI was assigned at least eight neighboring gold beads as fiducial markers, and for every ROI, Minflux imaging was performed for 1 h. Overall, the beads drift during the entire imaging duration was negligible with an average standard error less than 1 nm.

The optical setup contained three different illumination modalities provided through separate optical paths: (i) widefield excitation (488 nm, 560 nm and 640 nm), (ii) regularly focused excitation (560 nm or 642 nm) or focused activation (405 nm) and (iii) phase-modulated excitation (560 nm or 642 nm) leading to a 3D donut in the focal region. The scanning range of all beams was about 100 × 100µm^2^ in the lateral direction and 800 nm in the axial direction. A 1.4 numerical aperture oil immersion lens focused the excitation light into the sample and collected the fluorescence light. In contrast to the acquisition of fluorophore blinking events in regular SMLM, the transient binding of the imager strand with the docking strand acted as the blinking event. Imspector from Abberior Instruments was used as the microscopy acquisition software and images were rendered using Paraview software.

### Minflux image analysis

Minflux data were analyzed using a custom code written in Python. In Minflux, each “track” (tid) represents multiple detections (localizations) from single putative fluorophores. First, to filter spurious or background signals, we removed the tracks that contained less than four localizations. Next, to detect if each track is indeed corresponding to single fluorophores, a cluster analysis was performed using Gaussian mixture model on localizations that belonged to the same track to split distinct clusters and a new list of tracks (tid2) was created. The localizations belonged to each tid2 were combined into “events”, which provided the localization precision (Fig. 3C, standard error of the mean) of single fluorophores. Furthermore, another cluster analysis of the “events” was performed using DBSCAN for a range of epsilon values (10 to 100 nm) and minimum points, MinPts (1 to 3). The results were examined for the size and distribution of clusters, allowing selection of appropriate epsilon value for specific proteins. For VMAT2, an epsilon of 40 nm and MinPt 3 rendered a homogenous population of clusters with an average diameter of ∼47 nm, which is in the expected range for synaptic vesicles. The abnormally large-sized vesicles (>100 nm in diameter), though detected with the same settings, were not included in the average size determination for the vesicles but were separately compared between the *DJ-1* KO and control neurons (Fig. 5C).

### Protein extraction and western blot analysis

Cultured neurons were collected by scraping in chilled PBS and centrifuged at 500x g for 5 minutes. Cell pellets were successively homogenized using a lysis buffer containing 1% Triton™ X-100 (with 10% glycerol, 150 mM NaCl, 25 mM HEPES at pH 7.4, 1 mM EDTA, 1.5 mM MgCl2, and a proteinase inhibitor cocktail). Protein solution was quantified using a bicinchoninic acid (BCA) assay, and 10 µg were subjected to SDS-PAGE on gradient gels under reducing conditions. The gels were transferred onto polyvinylidene difluoride (PVDF) membranes and blocked with Licor Intercept blocking solution. The primary antibodies used for western blotting included: anti-GAPDH (1:10000; Millipore #MAB374; 1:10000; Cell Signaling #2118), anti-tubulin β-3 (1:10000; BioLegend #802001, #801201), anti-Tyrosine hydroxylase (1:1000; Millipore #AB152), anti-PARK7/DJ1 (1:5000, Abcam #ab76008), anti-α-synuclein (syn303; BioLegend; #824301; 1:200; C-20; Santa Cruz, #sc-7011-R; 1:1000), anti-Clathrin Heavy Chain (A-8) (Santa Cruz, #sc-271178; 1:100), anti-VMAT2 (abcam, #ab191121; 1:500). The blots were subsequently probed with the corresponding fluorescent secondary antibody (IRDye, Li-Cor) and visualized using the Odyssey Fc Imaging System (Li-Cor). For HRP-based blots, membranes were instead probed with secondary HRP coupled antibody (TrueBlot, Rockland), followed by revelation with Clarity Max Western ECL substrate (Biorad) and visualized using the Odyssey Fc Imaging System (Li-Cor).

### False fluorescent neurotransmitter assay

False fluorescent neurotransmitter 206 (FFN206; abcam, #ab144554) was used in accordance with the published in vitro assays(*36*). On days 25–30 of differentiation, 100,000 cells per well were seeded into 96-well plates (Falcon® 96-well Black/Clear Flat Bottom TC-treated Imaging Microplate) in preparation for live-cell staining. After that, the plates were cultured until day 90 of differentiation. After adding 100 μL/well of 1µM FFN206 diluted in culture medium, the whole plate was incubated for one hour at 37°C. The VMAT2 inhibitor tetrabenazine (Sigma-Aldrich, #T2952) functions as negative control of the assay. Neurons were treated with 10 µM tetrabenazine for 30 minutes before adding FFN206. Photometric analysis of fluorescence uptake was carried out utilizing excitation/emission wavelengths of 360–20/460–30 nm after one wash with PBS. Results were normalized to total protein concentration.

### Assessment of mitochondrial membrane potential using TMRE

In order to perform live-cell staining, 96-well plates (Falcon® 96-well Black/Clear Flat Bottom TC-treated Imaging Microplate) were seeded with 100,000 cells per well on days 25–30. The plates were then kept in culture until day 90. Unless otherwise noted, staining methods were carried out according to the manufacturer’s instructions. Two drops/mL of Hoechst 33342 (Invitrogen, #R37605) were used to stain the cell nuclei after TMRE (Abcam, #ab113852) had been diluted to 200 nM. After preparing dilutions in culture medium, 100 μL/well was added, and the mixture was incubated for 30 minutes. Following two PBS or 0.2% BSA washes, photometric analysis was performed using excitation/emission wavelengths of 360–20/460–30 nm (Hoechst 33342) and 540– 15/584–20 nm (TMRE).

### ATP assay

For ATP detection The ATP Colorimetric/Fluorometric Assay Kit (Sigma-Aldrich, #MAK190) was used. Freshly harvested cells were lysed in 100 μL of ATP Assay Buffer and deproteinized using a cellulose 10kDa MWCO spin filter (Sigma-Aldrich, #UFC501008). 2–50 μL of samples were added into duplicate wells of a 96 well plate and subsequently brought to a final volume of 50 μL by adding ATP Assay Buffer. Reaction Mixes were set up according to the schemes in ATP assay protocol from Sigma and 50 μL added to each of the wells. By gently pipetting up and down every well mix of components was ensured, followed by a 30 min incubation in the dark at room temperature protect the plate from light during the incubation. ATP standards for fluorometric detection were prepared as described in kit protocol. Fluorescent signal was measured by a BMG Labtech CLARIOstar Plus microplate reader using excitation/emission wavelengths of 535/587 nm.

### ATP treatment (incl. ATPyS non-active form)

For ATP treatment of dopaminergic neurons, 50 µM of either Adenosin-5′-triphosphat (ATP) (Sigma-Aldrich, #A6419) or its non-active form Adenosine-5’- O- (3-thiotriphosphate) (ATPγS) (BioLog, #A060) were applied to DJ-1-linked patient and *DJ-1* KO neurons for 72h prior to harvesting, refreshing the treatment and media every 24 hours.

### Transmission electron microscopy

iPSC-derived midbrain dopaminergic neurons were seeded onto plasma-treated ACLAR® (plastic) films (Science Services). We treated cells with prewarmed 2x fixative (5% glutaraldehyde (EM-grade, Science Services), 0.2M cacodylate buffer pH 7.4 (Science Services)) 1:1 to the culture medium. After 5 min this mixture was replaced by 1x fixative (2.5% glutaraldehyde in 0.1M cacodylate buffer) and incubated for 25 min on ice. Postfixation and -contrasting were performed by several washes in 0.1 M cacodylate buffer on ice followed by incubation in 1% osmium tetroxide (Science Services) and 0.8% potassium ferrocyanide (Sigma Aldrich) in 0.1 M sodium cacodylate buffer. Neuronal cells were further contrasted using 0.5% uranyl acetate in water (Science Services). After dehydration in ethanol and infiltration in LX112 resin (Ladd Research) the cell monolayer was embedded into resin blocks and cured for 48 hours at 60°C. Ultrathin sections (80 nm thick) were generated on a Leica UC6 ultramicrotome and adhered to formvar-coated copper grids (Plano). The sections were imaged on a JEM 1400plus (JEOL) transmission electron microscope using a XF416 camera (TVIPS) and the EM-Menu software (TVIPS). Image analysis was conducted using Fiji(*85*).

### Near infrared fluorescence (nIRF) detection of oxidized dopamine

nIRF assay for the detection of oxidized DA was performed as previously described in detail (*3*). Briefly, after being scraped in cold PBS, neurons were centrifuged for five minutes at 500x g. 1% Triton X-100 lysis buffer containing 10% glycerol, 150 mM NaCl, 25 mM Hepes pH7.4, 1 mM EDTA, 1.5 mM MgCl2 and proteinase inhibitor cocktail was used to homogenize the cell pellet. Boiling and sonication were used to recover insoluble pellets from a 100,000x g spin (30min, 4°C) in 2% SDS/50mM Tris pH 7.4, which were then incubated for 30 minutes at 4°C. After a 150,000x g spin, leftover pellets were further extracted in 1N NaOH and incubated over night at 55°C. Depending on the solubility of the remaining sample, the solution was either vortexed or sonicated to guarantee total solubilization. A lyophilization step using a Speed Vac Concentrator was done to lyophilize the fluid until a dried pellet remained. To remove remaining hydroxides and to lower pH levels, the pellet was first washed in Nanopure water. It was then lyophilized again until fully dry and then taken up in Nanopure water for final analysis. A stock of 10 mM oxidized DA was used to create the standard. To prepare oxidized DA, 10 mM DA and 20 mM NaIO4 were combined in D-PBS, vortexed, and then incubated for 5 minutes at room temperature. The solution was subsequently processed using centrifugation, sonication, and lyophilization techniques, same as the neuron samples. An Odyssey infrared imaging system (Li-Cor) with the 700 channel was used to scan the membranes after 5µl of each sample or standard dilution was dropped onto a Biodyne Nylon Transfer Membrane (Pall, #Pall-60209). Odyssey infrared imaging software was used to quantify the samples.

### Statistical analysis

Unpaired Student’s t-test was conducted for statistics in figs. 3A, D, F-G, 4A-B, 5C, S7E-F. ANOVA followed by Šidák’s multiple comparisons tests was conducted for statistics in figures 4D-F, 5D, 6A-B, 7A-E, S6A.

For the figs. 2A, 2B, and 2C, raw mass spectrometry data were processed using MaxQuant (v2.6.7.0), and statistical analysis was performed in Perseus (v2.1.3.0). Differences between groups were assessed using a two-sample Student’s t-test, with permutation-based FDR (False Discovery Rate) correction. Proteins were considered significantly differentially expressed if they had an FDR < 0.05 (–log₁₀(p) > 1.3) and a log₂ fold change > 1.

P-values <0.05 were considered significant, and all error bars in the figures represent standard error of the mean (SEM) (*P<0.05; **P<0.01, ***P<0.001). CTRL #1 and #2 values are combined as “CTRLs”, and *DJ-1* KO #1 and #2 values are combined as “*DJ-1*^KOs^”. A minimum of three differentiations were used for statistical analysis.

## Acknowledgments

We thank Rejko Krüger for providing iPSC lines from a DJ-1-linked PD patient and its CRISPR-edited control, Philip Seibler for providing iPSC lines from two healthy control individuals, and Dimitri Krainc for providing their respective CRISPR-edited *DJ-1* KO lines. We thank Christian Haass for providing equipment and assistance. We also thank Sabine Langer-Freitag and Iwona Sikora at the Institute of Human Genetics, Klinikum Rechts der Isar, School of Medicine, Technical University of Munich, for karyotyping of mouse iPSC lines.

## Author contributions

LFB conceived, designed and supervised this study. LMH, FG, MS, SFL, CB and SS designed the research. LMH, FG, AH, MR, AC-A, UM, SAM, SRN and LJ performed the experiments with support from LFB, MS, SFL, CB and SS. LMH, FG, AH, MR, AC-A, SAM, SRN, LJ and LFB analyzed the data. WW provided mouse embryonic fibroblasts. The manuscript was written by LMH, FG, AC-A, SS and LFB with contributions from all authors. All authors approved the manuscript.

## Data and materials availability

All data needed to evaluate the conclusions in the paper are present in the paper and/or the Supplementary Materials.

## Supplementary Materials

**Figure S1:**
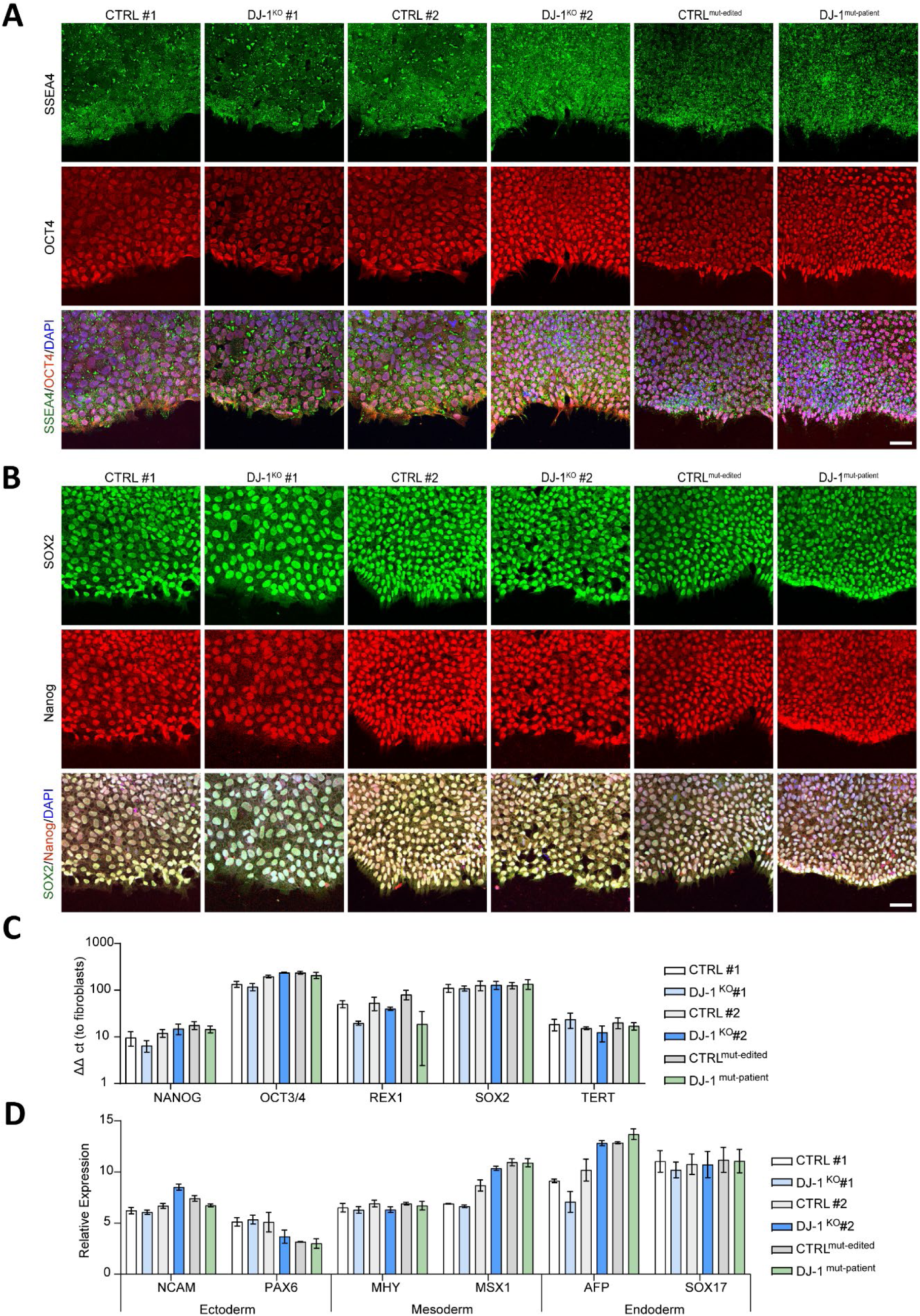
Characterization of iPSC lines from healthy controls and isogenic DJ-1 KOs, and DJ-1-linked PD patient and isogenic control. (**A-B**) Immunocytochemistry for pluripotency markers (A) OCT4 and SSEA4, or (B) NANOG and SOX2 in iPSCs from two healthy individuals (CTRL #1, CTRL #2) and from their isogenic DJ-1 KO lines (DJ-1^KO^ #1, DJ-1^KO^ #2), from a DJ-1-linked PD patient (DJ-1^mut-patient^) and its isogenic control (CTRL^mut-edited^). Merged presentation shows nuclear counterstaining with DAPI. Scale bar, 50μm. (**C**) Gene expression of NANOG, GDF3, OCT4, and SOX2 in iPSCs from two healthy individuals (CTRL #1, CTRL #2) and from their isogenic DJ-1 KO lines (DJ-1^KO^ #1, DJ-1^KO^ #2), from a DJ-1-linked PD patient (DJ-1^mut-patient^) and its isogenic control (CTRL^mut-edited^) relative to control fibroblasts (**D**) *In vitro* differentiation via embryoid body formation and qRT-PCR analyses of various differentiation markers for the three germ layers (endoderm: SOX17, AFP; mesoderm: MHY6, MSX1; ectoderm: PAX6, NCAM) in iPSCs from two healthy individuals (CTRL #1, CTRL #2) and from their isogenic DJ-1 KO lines (DJ-1^KO^ #1, DJ-1^KO^ #2), from a DJ-1-linked PD patient (DJ-1^mut-patient^) and its isogenic control (CTRL^mut-edited^) that were differentiated for 14 days.

**Figure S2:**
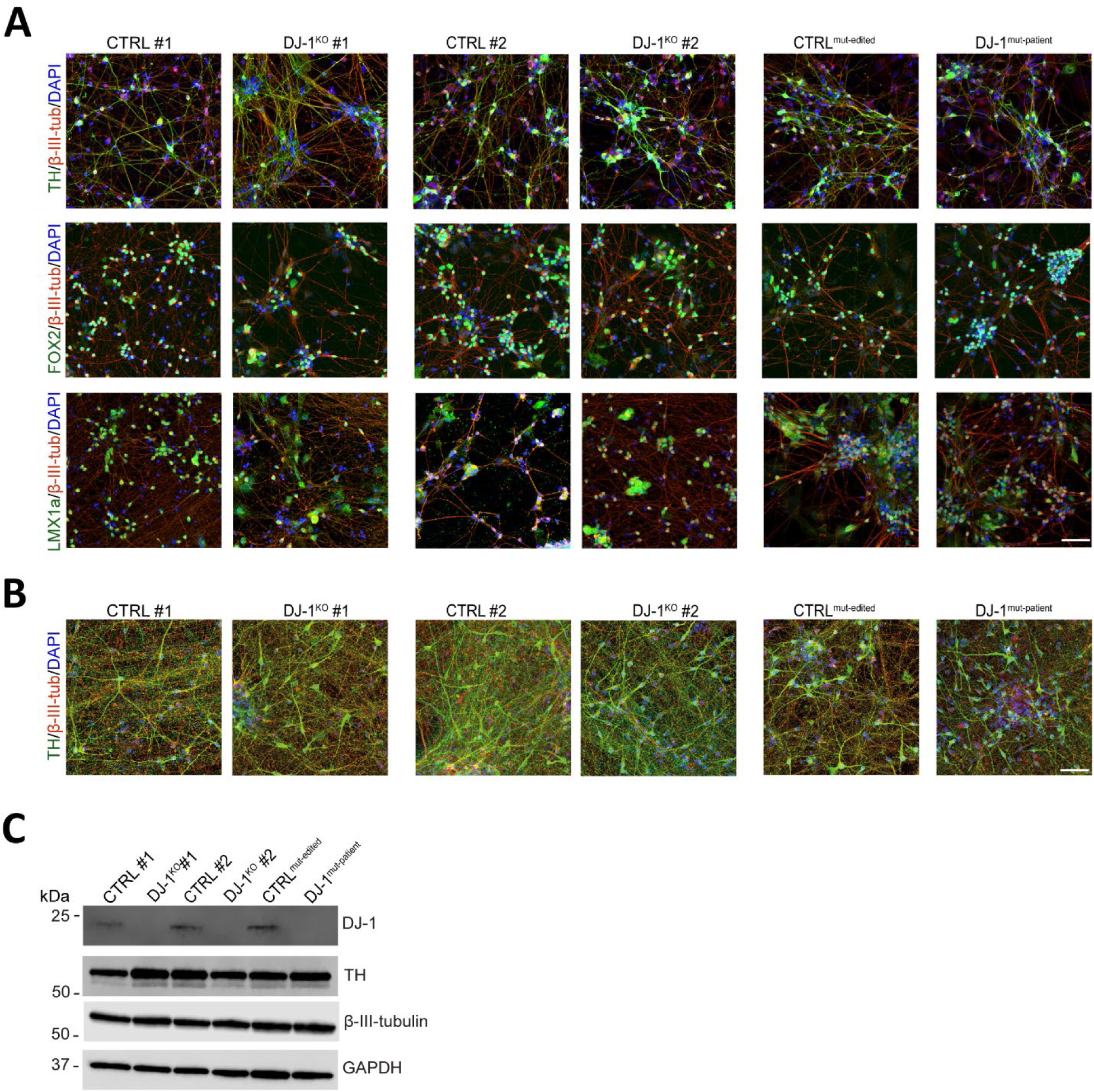
**Characterization of differentiated dopaminergic neurons from healthy controls and isogenic DJ-1 KOs, and DJ-1-linked PD patient and isogenic control.** (**A**) Immunocytochemistry of midbrain, dopaminergic and neuronal markers in iPSC-derived dopaminergic neurons from two healthy individuals (CTRL #1, CTRL #2) and from their isogenic DJ-1 KO (DJ-1^KO^ #1, DJ-1^KO^ #2) counterparts, from a DJ-1-linked PD patient ((DJ-1^mut-patient^) and its isogenic control (CTRL^mut-edited^) at day 30 of differentiation. Antibodies against FOXA2, LMX1a, TH and β-III-tubulin were used. Scale bar, 50μm. (**B**) Visualization of TH expression by immunocytochemistry in iPSC-derived dopaminergic neurons from two healthy individuals (CTRL #1, CTRL #2) and from their isogenic DJ-1 KO (DJ-1^KO^ #1, DJ-1^KO^ #2) counterparts, from a DJ-1-linked PD patient ((DJ-1^mut-patient^) and its isogenic control (CTRL^mut-edited^) at day 70 of differentiation, and quantification. Antibodies against TH and β-III-tubulin were used. Scale bar, 50μm. (**C**) Representative immunoblot of TH and β-III-tubulin levels in T-soluble neuronal lysates from two healthy individuals (CTRL #1, CTRL #2) and from their isogenic DJ-1 KO (DJ-1^KO^ #1, DJ-1^KO^ #2), from a DJ-1-linked PD patient ((DJ-1^mut-patient^) and its isogenic control (CTRL^mut-edited^) at day 70 of differentiation. Immunoblot analysis of DJ-1 confirms lack of protein expression in DJ-1 KO and DJ-1-linked PD patient lines. GAPDH was used as a loading control.

**Figure S3:**
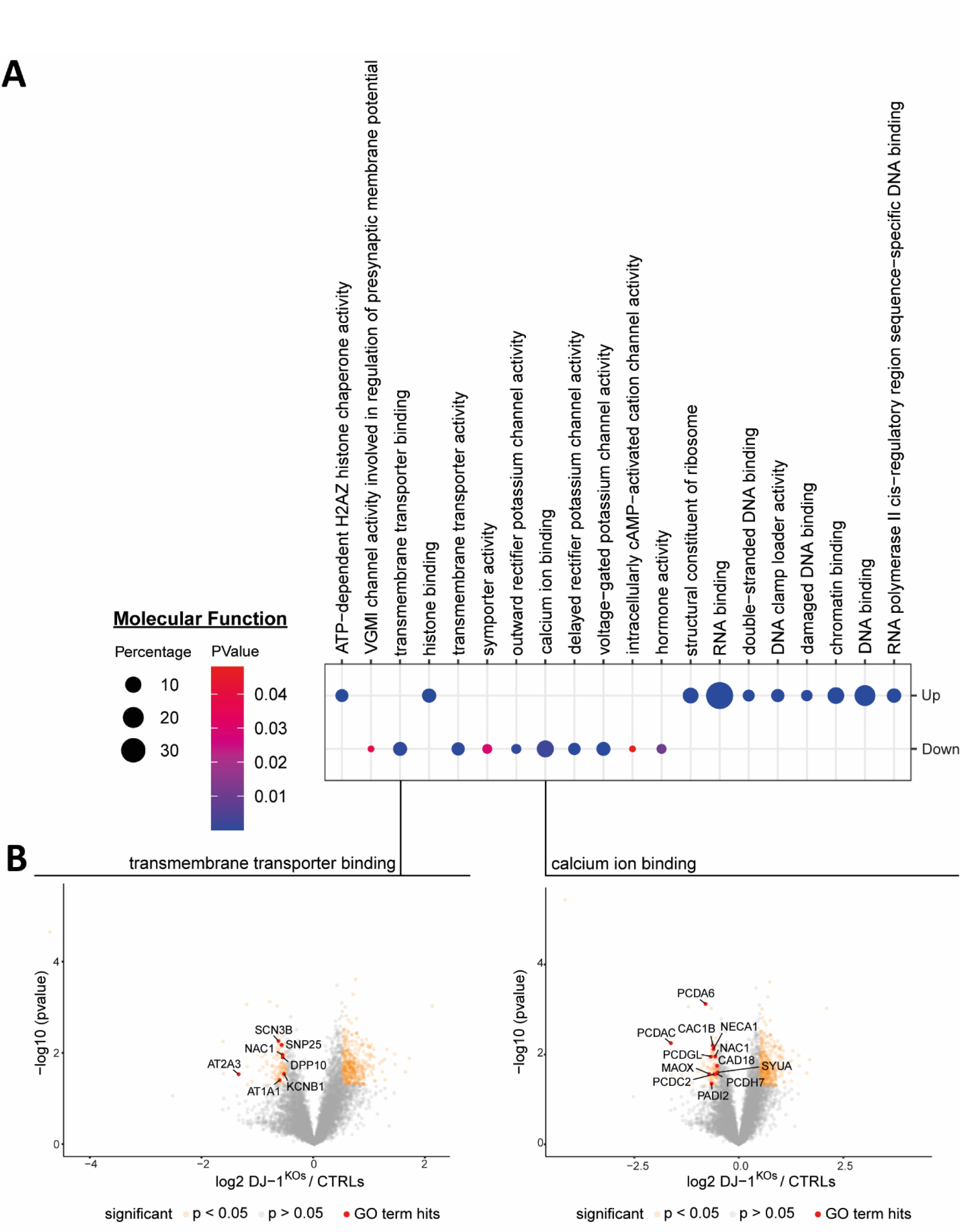
Gene ontology enrichment analysis for molecular function of DJ-1-deficient dopaminergic neurons. (**A**) Results of gene ontology (GO) enrichment analysis for molecular function (GOTERM_MF_DIRECT) using the web tool DAVID (v 6.8). Proteins with a consistently increased or reduced abundance were compared to the background of all proteins quantified for both comparisons. The dot plots show the total gene number of a term in percentage as dot size and the p-values as a color gradient. The top 10 terms for either enriched (Up) or downregulated (Down) proteins are presented. (**B**) Volcano plot of DEGs from both *DJ-1* KO lines versus their isogenic control lines for prominent enrichment of downregulated proteins for the GO terms “transmembrane transporter binding” and “calcium ion binding” associated with molecular function.

**Figure S4:**
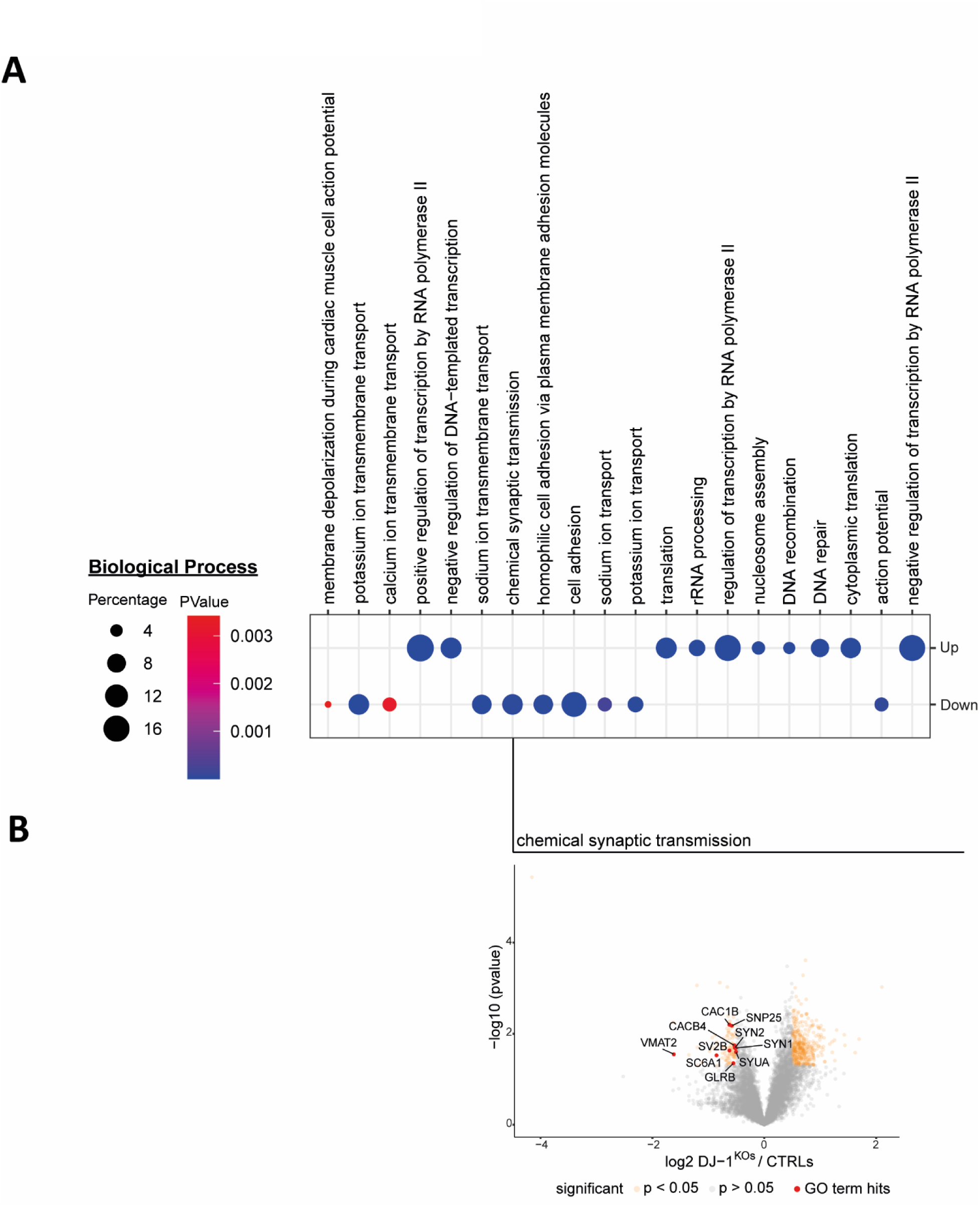
Gene ontology enrichment analysis for biological process of DJ-1-deficient dopaminergic neurons. (**A**) Results of gene ontology (GO) enrichment analysis for biological process (GOTERM_BP_DIRECT) using the web tool DAVID (v 6.8). Proteins with a consistently increased or reduced abundance were compared to the background of all proteins quantified for both comparisons. The dot plots show the total gene number of a term in percentage as dot size and the p-values as a color gradient. The top 10 terms for either enriched (Up) or downregulated (Down) proteins are presented (**B**) Volcano plot of DEGs from both DJ-1 KO lines versus their isogenic control lines for prominent enrichment of downregulated proteins for the GO term “chemical synaptic transmission” associated with biological process.

**Figure S5:**
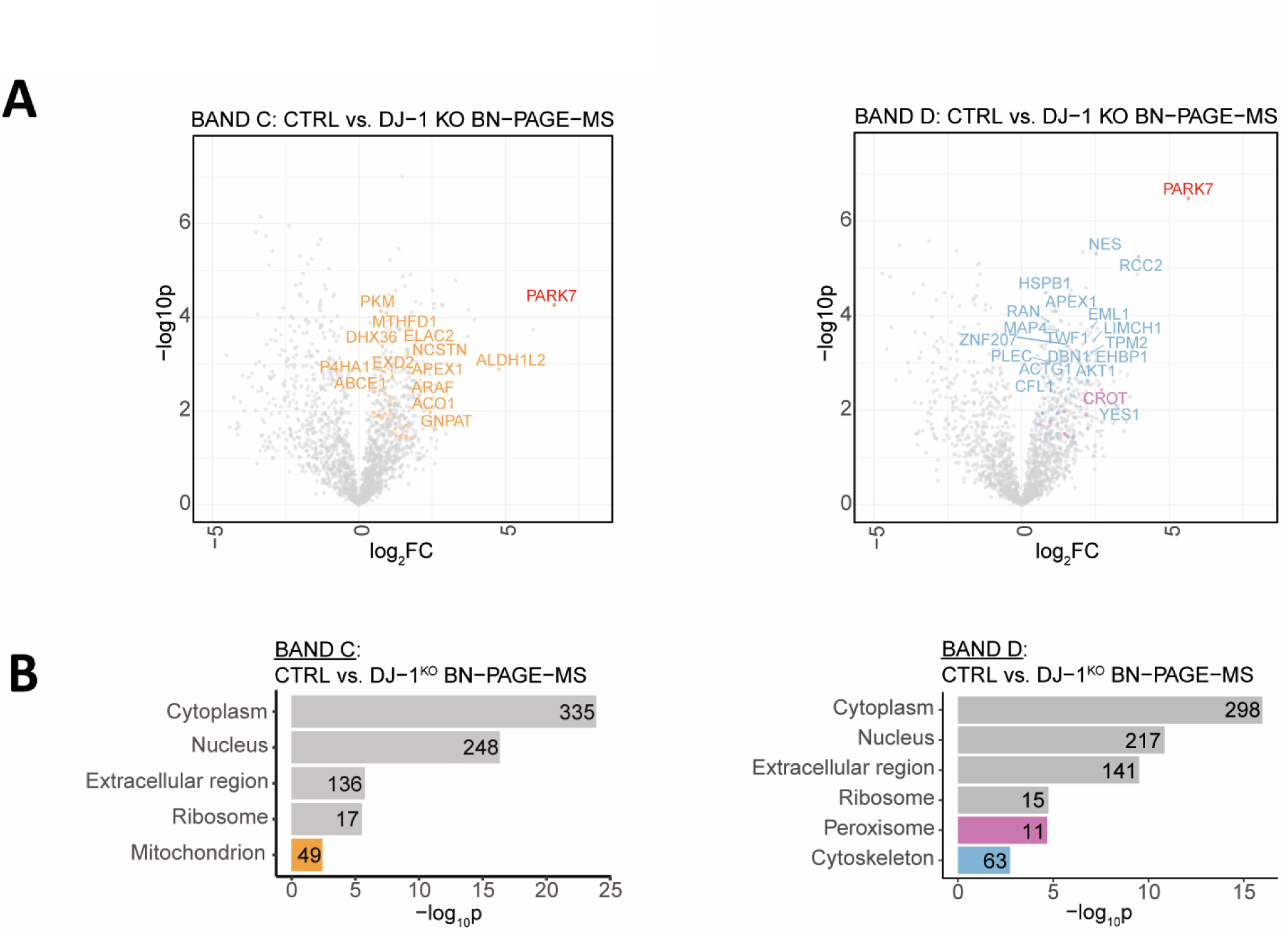
Volcano plots and GO cellular component analysis of DJ-1 co-migrants in bands “C” and “D”. (**A**) Volcano plots of the proteins in bands “C” and “D” co-migrating with DJ-1, identified by comparing control to *DJ-1* KO neurons (two-sided t-test, n = 4 independent experiments). Highlighted are representative DJ-1 co-migrants associated with the mitochondria (orange), peroxisome (pink), and the cytoskeleton (blue). PARK7 is highlighted in red. (**B**) GO cellular component analysis, using SubcellulaRVis, of DJ-1 co-migrants in bands “C” and “D” (log2FC > 0.5, p < 0.05).

**Figure S6:**
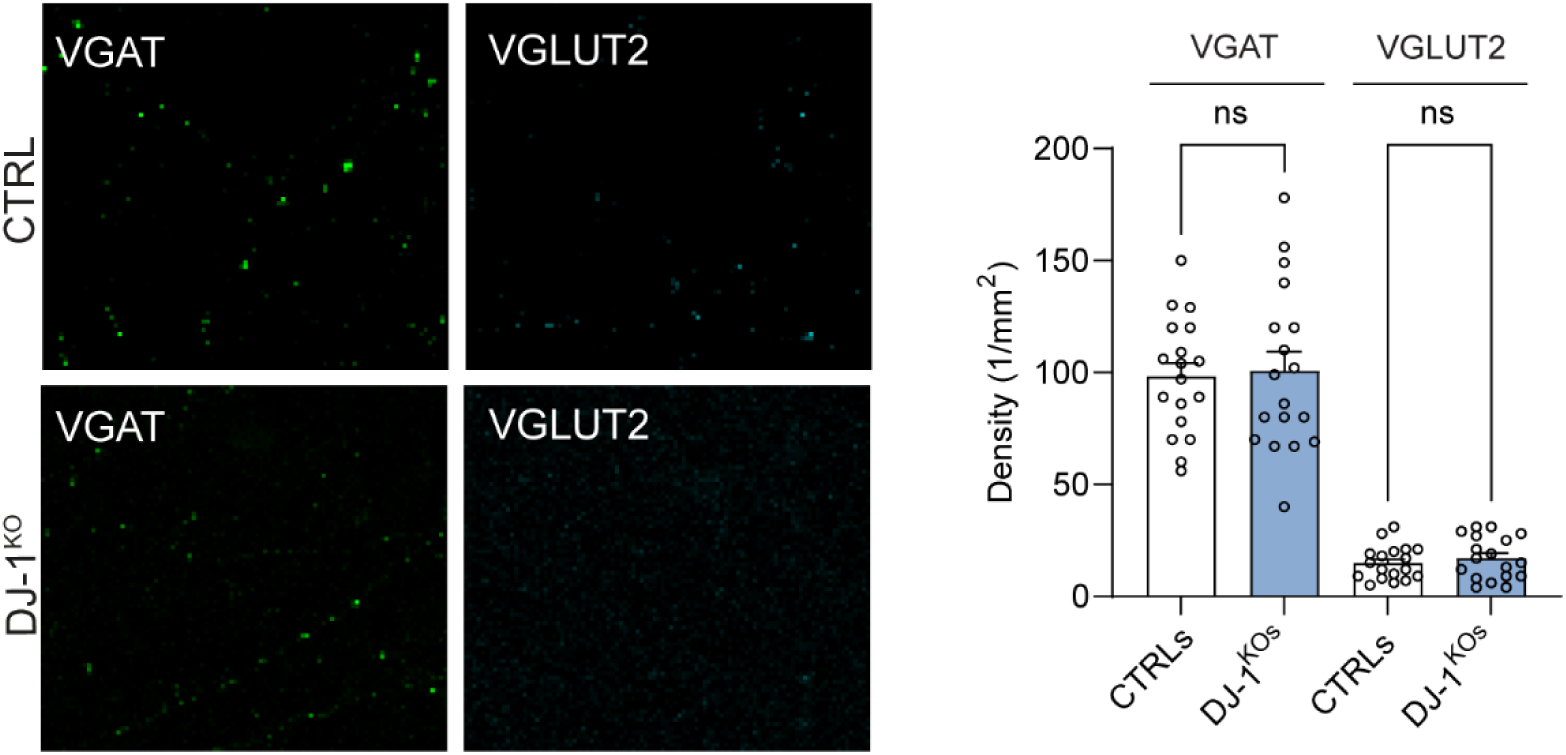
Analysis of VGAT- or VGLUT2-positive synapses in control and DJ-1 KO neurons. Representative confocal images of VGAT- or VGLUT2-positive synapses in control (top) and *DJ-1* KO neurons (bottom). Individual synapses were visualized by staining VGAT or VGLUT2, and quantified. Quantification shows no difference in density of VGAT- or VGLUT2-positive synapses in *DJ-1* KO neurons compared to controls. Scale bar, 20µm (n=18).

**Figure S7:**
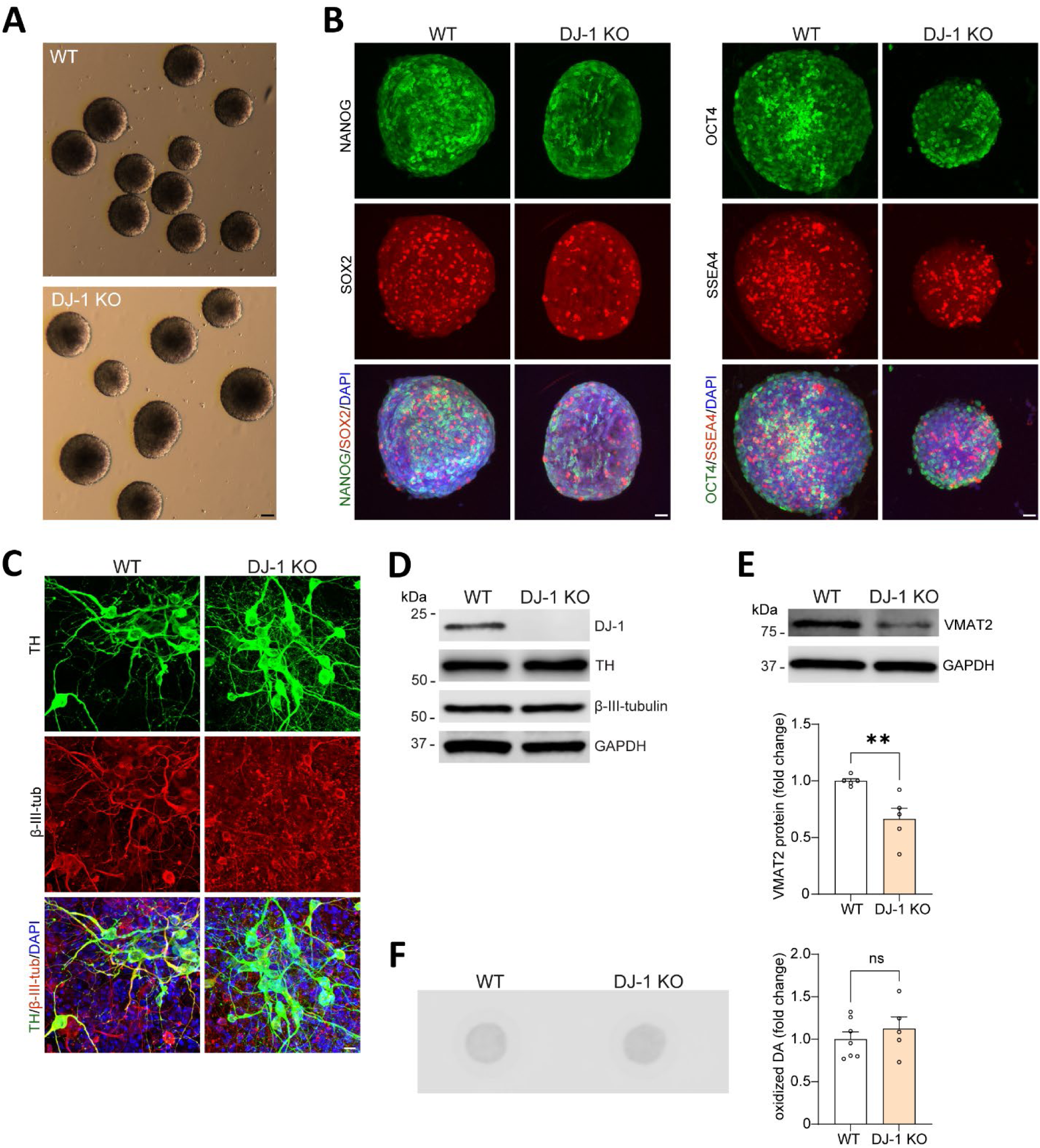
Characterization of WT and *DJ-1* KO mouse iPSC lines and of iPSC-derived WT and *DJ-1* KO mouse dopaminergic neurons. (**A**) Representative brightfield image from WT (top) and *DJ-1* KO (bottom) iPSC spheres. Scale bar, 100μm. (**B**) Immunocytochemistry for pluripotency markers NANOG and SOX2 (left), or OCT4 and SSEA4 (right) in iPSCs from WT and *DJ-1* KO mouse. Merged presentation shows nuclear counterstaining with DAPI. Scale bar, 25μm. (**C**) Immunocytochemistry of the dopaminergic neuron marker TH and the neuronal marker β-III-tubulin in iPSC-derived dopaminergic neurons from WT and *DJ-1* KO mouse iPSCs at day 30 of differentiation. Merged presentation shows nuclear counterstaining with DAPI. Scale bar, 10μm. (**D**) Representative immunoblot of TH and β-III-tubulin levels in T-soluble neuronal lysates from WT and *DJ-1* KO iPSC-derived mouse dopaminergic neurons at day 30 of differentiation. Immunoblot analysis of DJ-1 confirms lack of protein expression in *DJ-1* KO line. GAPDH was used as a loading control. (**E**) Immunoblot analysis of VMAT2 in T-soluble neuronal lysates from WT and *DJ-1* KO mouse dopaminergic neurons (n = 5). GAPDH was used as loading control. (**F**) Representative image of oxidized DA by nIRF in WT and *DJ-1* KO mouse dopaminergic neurons, and quantification (n = 5-7).

**Table S1. Whole-cell proteomics in *DJ-1* KO versus isogenic control iPSC-derived dopaminergic neurons.**

* see attached Excel file “Table S1”

**Table S2. Protein group identification and quantification data (MaxQuant) for the BN-PAGE co-migration experiment performed on *DJ-1* KO and isogenic control iPSC-derived dopaminergic neurons.**

* see attached Excel file “Table S2”

**Table S3. Protein group identification and quantification data (MaxQuant) for the DJ-1 IP-MS experiment performed on *DJ-1* KO and isogenic control iPSC-derived dopaminergic neurons.**

* see attached Excel file “Table S3”

